# A Family VIII Esterase with Dual Activities: Bioplastic Depolymerization and Beta-Lactam Antibiotics Hydrolysis

**DOI:** 10.1101/2025.09.01.670660

**Authors:** Harry Lerner, Nele Charlott Meier, Léa Bernabeu, Marcel Eck, Stefan Mecking, David Schleheck

**Affiliations:** Department of Biology, University of Konstanz, Konstanz, Germany; Department of Chemistry, University of Konstanz, Konstanz, Germany; Limnological Institute, University of Konstanz, Konstanz, Germany *

## Abstract

Microorganisms in the plastisphere are associated with plastic degradation, as well as with an unusually high occurrence of antibiotic resistance genes (ARGs), suggesting plastic debris as a potential vector for antibiotic-resistant microorganisms. In this study, we investigated microbial communities associated with the degradation of aliphatic long-chain polyester (LCAP) bioplastics in forest soil. Sequencing analysis revealed a family VIII esterase with structural similarity to type C β-lactamases. Structural modeling and substrate docking analysis indicated catalytically favorable binding of both LCAP and β-lactam antibiotics. Heterologous expression and *in-vitro* activity testing confirmed its dual functionality as plastic depolymerase and β-lactam hydrolase. Sequence-based predictions identify the enzyme as a membrane-associated lipoprotein, with suggested further secretion via outer-membrane vesicles (OMVs), offering potential ecological benefits in competitive plastisphere environments. These findings highlight an enzyme with a rare substrate spectrum, bridging plastic and antibiotics degradation and suggesting an intriguing biochemical connection that warrants further investigation of microbial evolution in the plastisphere and its potential implications for the spread of antibiotic resistance.

## Introduction

Plastic waste and its deterioration into micro- and nanoplastics has developed into one of the major environmental and health concerns of our time.^1–4^ In addition to being physical pollutants, plastic debris is also considered a vector for chemical contaminants and microbial colonization.^5–8^ Microorganisms inhabiting the plastisphere, i.e., the biofilm community on plastic surfaces, have been shown to harbor disproportionately high levels of antibiotic resistance genes (ARGs), raising concerns about plastic-mediated dissemination of ARGs throughout terrestrial and aquatic ecosystems.^9–13^ This enrichment of ARGs has largely been attributed to the increased occurrence of horizontal gene transfer (HGT) within plastisphere communities, although no clear fitness driving factor has been reported.

One driving factor of selection for resistance may be due to plastic associated contaminants.^10,12^ Microplastics are known to form ‘eco-coronas’, i.e., layers of environmental (bio)molecules, including heavy metals, pharmaceuticals, and antibiotics, that adsorb to their hydrophobic surface.^5–7^ Interestingly, a recent study assessed a higher antimicrobial resistance (AMR) risk for biodegradable microplastics compared to nondegradable ones and reported the genetic co-localization of degradation genes, ARGs, and virulence factors.^14^ As such, the plastisphere, particularly of biodegradable plastics, may exert selection pressure for resistance, while the plastic itself presents a potential carbon and energy source for microorganisms capable of depolymerizing the materials into water-soluble monomers (or oligomers) that can enter central metabolism. Thus, organisms capable of surviving in the face of antibiotics, and simultaneously accessing plastic-derived carbon for growth, would attain a pronounced fitness advantage within the plastisphere.

However, the utilization of plastic as a growth substrate is biochemically and ecologically challenging. Plastics are nutrient-poor ‘carbon deserts‘ that require the costly synthesis and secretion of extracellular depolymerizing enzymes, before a further intracellular utilization of the monomers is possible. This process catalyzed within a biofilm community carries the pronounced risk of loss of the monomers to commensal, and thus competing microorganisms. Nonetheless, the vast majority of plastic-degrading enzymes known to date appear to be freely secreted enzymes, making their host organisms particularly vulnerable to exploitation by non-producers (cheaters), as well as potentially inefficient, especially in aquatic environments, where both the secreted enzymes and hydrolysis products may diffuse away.

Recent studies have identified a membrane-associated PET-degrading esterase in *Rhodococcus pyridinivorans*, as well as the involvement of outer membrane vesicles (OMVs) in polyurethane degradation, suggesting a strategic shift towards retaining plastic depolymerization activity at or near the cell surface.^15,16^ This mechanism would help degraders retain access to the monomers they liberate, representing a logical advantage in highly competitive plastisphere communities. As such, despite hundreds of reported (often putative) plastic depolymerizing enzymes recorded in databases,^17,18^ their diversity and specialization likely remains underrepresented, as microorganisms continue to adapt to this new type of substrate. This is further evidenced by the recently reported functional overlap between plastic depolymerization and β-lactam antibiotic degradation for a metagenomically sourced esterase that can hydrolyze PET,^19^ suggesting a potential link between polymer degradation and β-lactam resistance in the plastisphere beyond HGT. While this functional overlap appears rare, it may indicate the co-selection of polymer degradation and antibiotic resistance in the plastisphere, where selective pressures may benefit such dual functionality.

Particularly family VIII type esterases hold a high potential for catalytic activity toward β-lactam antibiotics. This enzyme family contains catalytic motives typical for both enzyme groups along with additional structural similarities to class C β-lactamases. Despite this, the majority of family VIII esterases are incapable of β-lactam hydrolysis,^20^ and few have been reported to depolymerize plastic polymers.^21–23^ However in recent years, reports of family VIII esterases with β-lactamase activity have increased, including activity towards third- and fourth-generation cephalosporins, classified as by the WHO as critically important antimicrobials.^24–30^ This suggests a yet under-investigated contribution of this enzyme family towards antimicrobial resistance in natural environments.

In this study, we investigated the biodegradation of an emerging type of bioplastic, long-chain aliphatic polyesters (LCAPs), in forest soil using field trials and laboratory microcosms.

LCAPs are chemically recyclable and biodegradable plastics comprised of ester-linked long-chain (e.g., C18 alkyl) dialcohol and dicarboxylic acid monomers that can be biologically sourced from plant oils.^31–33^ Other than biodegradable short-chain aliphatic polyesters such as polycaprolactone (PCL), they can exhibit material properties similar to high-density polyethylene (HDPE),^31^ making them sustainable alternatives to conventional polyolefins.

LCAP degradation was assessed via electron microscopy and the tracking of substrate induced CO_2_ evolution for the field trial and microcosm set-up, respectively. To identify organisms and functions potentially involved in LCAP degradation, we performed metagenomic and amplicon-based sequencing of microbial communities in the microcosm set-up during active degradation. The bioinformatic analysis allowed for the identification of candidate enzymes with the predicted capacity to hydrolyze LCAPs, while at the same time matching the taxonomy of taxa enriched in the LCAP-treated microcosms. Among the candidate enzymes, we identified a previously undescribed family VIII esterase, which we designate LCPH1 (Long-Chain Polyester Hydrolase 1). LCPH1 showed similarities to both α/β-fold hydrolases and type C β-lactamases, and *in-silico* docking analysis suggested its potential for a rare dual functionality through favorable binding of β-lactam antibiotics as well as polyester substrates. This predicted substrate range was confirmed through heterologous expression, purification and *in-vitro* testing of the enzyme activity, demonstrating both the depolymerization of LCAP, as well as the hydrolysis of the β-lactam antibiotics Penicillin G and Ampicillin. Notably, LCPH1 is predicted to be a membrane-anchored lipoprotein, raising the possibility of surface localization and OMV-mediated secretion, which may provide a selective advantage for nutrient retention in the plastisphere environment. Hence, this study reports the first instance of a polyester plastic-depolymerizing family VIII esterase with confirmed β-lactamase activity, highlighting a functional link between plastic degradation and antibiotic resistance and providing new insights into the ecology of plastic degradation in soil environments.

## Results

### LCAPs show signs of *in-situ* degradation and are mineralized in forest-soil microcosms

Films of LCAP plastics PE-12,12 and PE-18,18 were examined for biodegradation in the humus layer of a pristine mixed forest soil for nearly 15 months (Fig. S1). The retrieved plastic films were macroscopically intact, but appeared more turbid and showed signs of breakage or embrittlement. Their examination by scanning electron microscopy (SEM) revealed substantial surface deterioration and material loss in form of frequent and clustered, approximately bacterial-cell sized, often rod-shaped cavities in the film surfaces (Fig. 1 & Fig. S2). The observed surface alterations were attributed to microbial degradation. Incomplete degradation was likely due to the weather conditions during the incubation period across the summers 2021 to 2022, which were marked by heat waves and very low precipitation.

**Figure 1.**
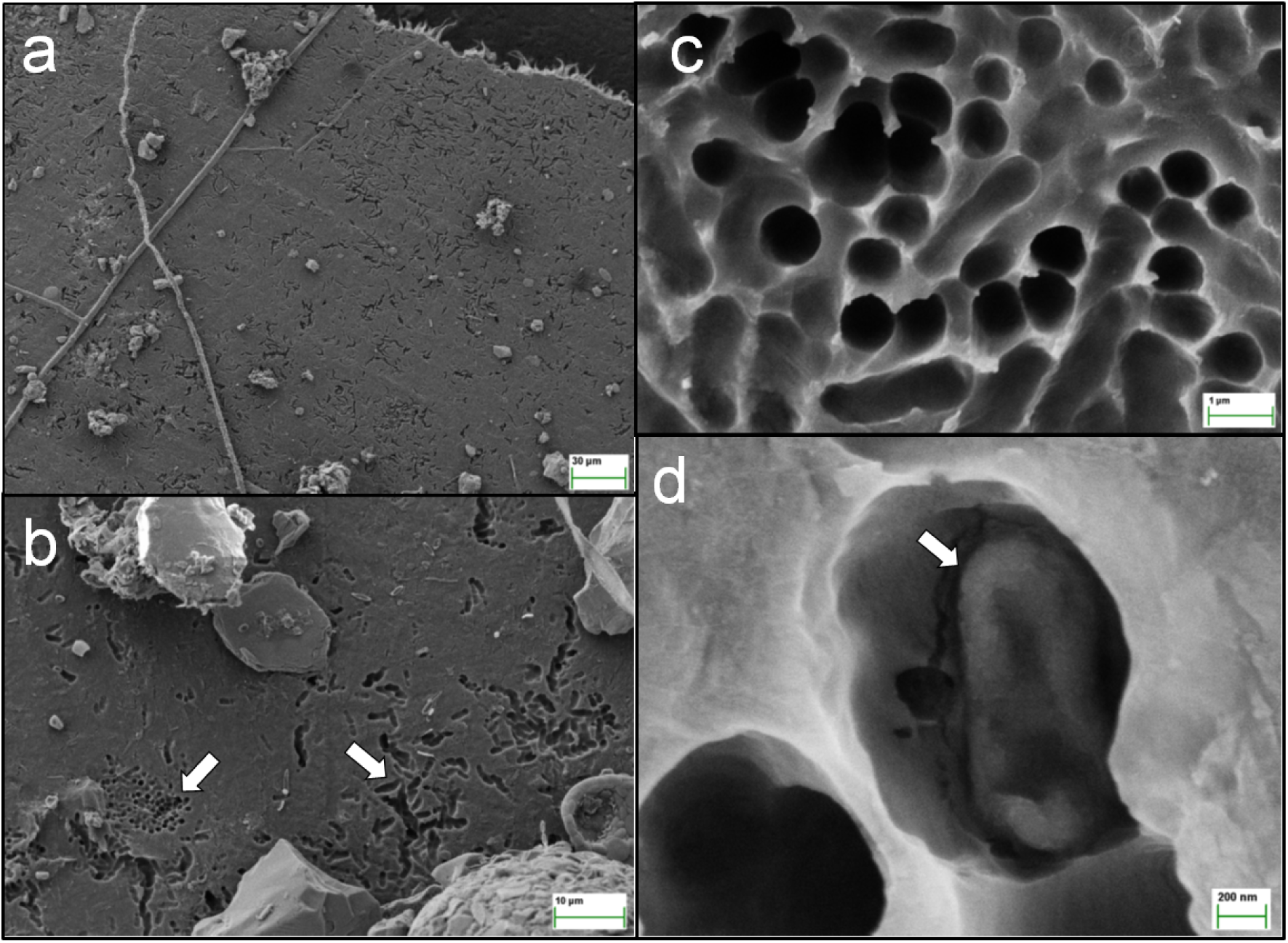
Scanning electron microscopy (SEM) images of LCAP films incubated undisturbed for more than a year in pristine forest soil. Representative images at increasing magnification are shown (see scale bars in panels a - d). The structural damage on the bioplastic films after incubation, which were absent from non-incubated films (see Supplementary information, Fig. S2), obviously resulted from a very localized degradation of the plastic material, visible in form of microbial-cell sized cavities (a). The two arrows in panel b point at two different clusters of microbial-cell sized cavities with different shapes and sizes. A representative cluster of microbial-cell shaped cavities is shown in higher magnification in c. The arrow in panel d points at an object inside of such a cavity that might represent a (damaged) microbial cell, however, these were found very rarely; empty cavities were more representative (a - c).

To assess the LCAP-degradation potential in forest soil under more controlled conditions, soil was collected close to the location of the *in-situ* incubation and used to set up microcosms. The soil was spiked with powders of LCAP plastics PE-2,18, PE-18,18 or PE-12,12, as well as of three reference materials, poly(hydroxybutyrate-co-hydroxyvalerate) (PHBV), polycaprolactone (PCL) and HDPE. Cellulose was used as an additional control. Substrate-induced CO_2_ evolution was quantified over up to 318 days of incubation in reference to a set of non-spiked control microcosms.

During the initial 23 days, insufficient sampling frequency and capacity of the CO_2_ traps resulted in non-trapped CO_2_, reducing the accuracy of measurements during that phase. While this affected all samples, the impact was likely highest for cellulose-spiked samples, where degradation started within few days (Fig. 2). The capacity of the CO_2_ traps was increased by using a higher NaOH concentration for the remainder of the experiment (see arrow in Fig. 2).

**Figure 2.**
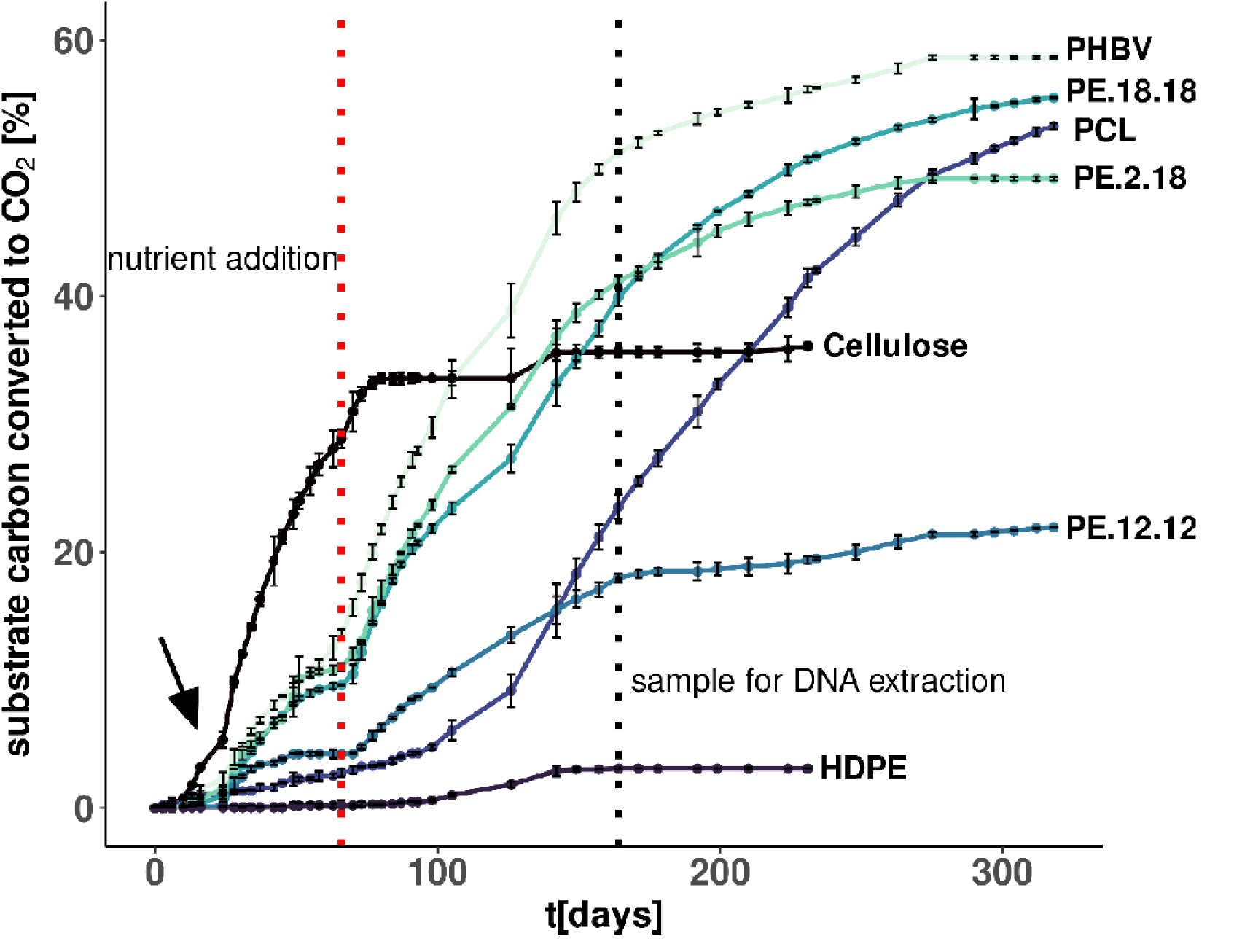
Substrate-specific respiration of forest-soil microcosms supplemented with powders of polymers PE-12,12, PE-18,18 and PE-18,2 in comparison to control polymers. Cumulative CO_2_ production is shown in percent of total polymer-carbon released as CO_2_, each after subtracting the basal CO_2_ production of untreated soil microcosms (blanks). The experiment was conducted using triplicate incubations for each polymer tested; standard deviation is indicated by error bars. The time point of external nutrient addition to the microcosms (see main text; N, P and trace metals) is marked by a red dotted line (day 66). The time point of sampling of all soil microcosms for total-DNA extraction and microbial community analysis by next generation sequencing, is indicated by the black dotted line (day 164). Control polymers used were poly(hydroxybutyrate-co-hydroxyvalerate) (PHBV), polycaprolactone (PCL) and high-density polyethylene (HDPE), as well as cellulose. The arrow indicates the time point after which the capacity of the CO_2_ traps had been increased (see main text).

After 77 days, CO_2_ production for cellulose plateaued at ∼36% of cellulose-carbon converted to CO_2_. This value is lower than the expected 50% observed during aerobic growth, where typically half the substrate carbon is respired as CO₂ and the other half is incorporated into microbial biomass.^34^ The lower recovery is likely due to unmeasured (non-trapped) CO during the early incubation phase. In contrast, HDPE, representing nondegradable plastics, displayed a cumulative CO_2_ production of only 3% of substrate carbon after a total incubation time of 231 days (Fig. 2).

After 25 days, CO₂ production increased for PE-18,18, PE-2,18, PHBV, and, to a lesser extent, PE-12,12 and PCL (Fig. 2). By day 50, respiration rates plateaued, with cumulative CO₂ accounting for only ∼10% of substrate-carbon for PHBV, PE-18,18, and PE-2,18, and <5% for PCL and PE-12,12. We considered that this plateau likely reflected nutrient limitation, as indicated by our soil’s low pH (4.3), which may have affected nutrient bioavailability despite a favorable C:N ratio (Table S2). Accordingly, microcosms were supplemented with a mineral-salts mixture (including ammonium, phosphate, and trace metals) at day 66 (Fig. 2, red dashed line).

Upon nutrient addition, accelerated respiration was observed within few days, most pronounced for PHBV, PE-18,18 and PE-2,18, and to a lower extent also for PE-12,12 and cellulose (Fig. 2). In contrast, the CO_2_ production for HDPE and PCL exhibited no immediate increase. Notably, the nutrient amendment did not stimulate background soil respiration (Fig. S3). On day 164 of the incubation, sub-samples were retrieved from all soil microcosms for DNA extraction and sequencing-based microbial community analysis (black dashed line in Fig. 2). The remaining soil was incubated further and cumulative CO_2_ production was recorded.

After ∼250 days, the cumulative CO_2_ production for PE-18,18 and PE-2,18, as well as for PHBV, plateaued between 49% and 58%, suggesting near-complete degradation. For PE-12,12, the cumulative CO_2_ production plateaued at only ∼22% (see Discussion). For PCL, mineralization reached ∼53% by day 318, indicating delayed but likely complete degradation.

### Phylogenetic analysis of bacterial and fungal communities in polymer-degrading soil microcosms

Substrate-specific changes in bacterial and fungal community compositions were assessed by amplicon-based sequencing analysis of DNA isolated after 164 days. Principal coordinates analysis (PCoA) and PERMANOVA (p = 0.001) revealed clear clustering of bacterial and fungal communities corresponding to the respective polymer treatment (Fig. 3a). The bacterial communities in the soil degrading PE-12,12, PE-18,18 and PE-2,18 were most dissimilar compared to non-spiked soil, and clustered closely together regardless of the LCAP variant used; only replicate 2 of the PE-18,18 treatment grouped closer to the cellulose cluster. The PHBV and cellulose replicates grouped close to the 0-point of the PCoA, and the PCL replicates grouped relatively close to the non-spiked controls. HDPE-treated soils also clustered near the non-spiked soil, although replicates were relatively dispersed on axis 2 (Fig. 3a). Similar clustering was observed for the ITS amplicon sequencing data, with LCAP-treated samples showing the highest degree of dissimilarity to the non-spiked soil (Fig 3a).

**Figure 3.**
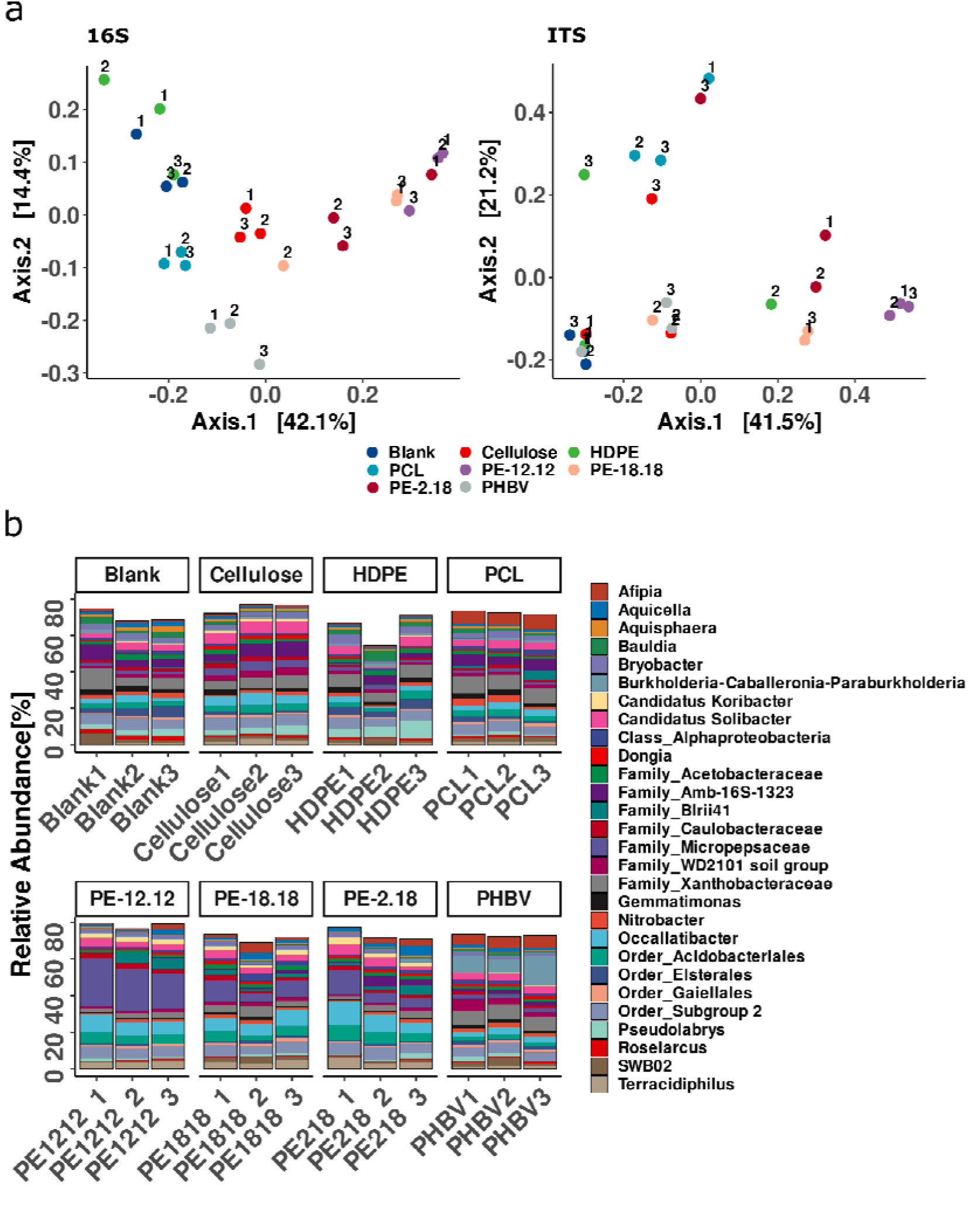
Principal Coordinates Analysis (PCoA) of 16S rRNA-gene and ITS amplicon sequencing data (a) and bar plots illustrating the 16S rRNA gene diversity (genus level) of the bacterial communities in soil samples incubated with the different polymers (b). The PCoA is based on Bray Curtis dissimilarity metrics. Treatments are indicated in different colors and individual replicates are numbered. Percentages given at the axes indicate the dissimilarity explained by the respective axis. **(b)** The community composition is shown for all replicate treatments. ASVs assigned to the same genus were combined. When a genus could not be determined with sufficient confidence, the last taxonomic level assigned is indicated. Only ASV groups with a mean relative abundance of >1% are depicted.

Bar graphs at the genus level further illustrate the marked relative-abundance change of bacterial ASVs in the polymer-treated microcosms (Fig. 3b). The LCAP-treated microcosms showed the evident increase of ASVs assigned to family Micropepsaceae, as well as of the closely related Acidobacteriaceae genera *Terracidiphilus* and *Occallatibacter*, while the PHBV treatments showed a distinct increase of ASVs assigned to genera of the BCP group (*Burkholderia*, *Caballeronia*, and *Paraburkholderia*) (Fig. 3b). The genus *Afipia* appeared in increased abundance especially in PCL- and PHBV-treated microcosms, but also in four of the LCAP-treated replicates. For the ITS-based fungal community, the bar graph showed a marked reduction of the GS11 clade compared to the non-spiked samples and an increased dominance of an ASV group with unidentified taxonomic affiliation (*incertae sedis*) in several LCAP-treated samples (Fig. S4a). Two PE-2,18 replicates showed an additional increase in sequences belonging to the *Rozellomycota*, which likewise showed a strong increase in one cellulose replicate and two PCL replicates (Fig. S4a). Overall, the abundance changes in the fungal community appeared to be considerably lower compared to the observed changes in the bacterial communities, though few dominant groups with poor taxonomic resolution differed strongly between treatments and replicates (Fig. S4a).

A differential abundance analysis based on the log2-fold change (LFC) was used to identify those taxa that significantly increased in the polymer treatments. To this end, non-spiked controls and the inactive HDPE-treated microcosms were grouped together to serve as the reference group. Likewise, microcosms treated with the three types of LCAP were grouped together, while the replicates treated with cellulose, PHBV and PCL formed separate groups.

The analysis conducted on the ITS dataset resulted in the detection of significantly different abundances for 10 ASVs (9 increased, 1 decreased) in the LCAP group, all of which provided poor taxonomic information beyond the fungal kingdom (*incertae sedis*). For the PCL group, two ASVs increased significantly, both of which belonged to the Phylum *Rozellomycota* (Fig. S4b).

The LCF analysis of the 16S rRNA-gene dataset resulted in the detection of many differentially abundant taxa and was therefore conducted at the genus and ASV level (Fig. 4ab). Most differences were detected in the LCAP group, followed by the cellulose and the PHBV groups, while only sequences associated with *Afipia* were significantly increased in the PCL group (Fig. 4a). Seven taxa were uniquely increased in the LCAP group: the genera *Mucilaginibacter* (Sphingobacteriia) and *Gemmata* (Planctomycetia), the non-cultivated taxa KF-JG30-25 (Acidobacteriia), Blrii41 (Acidobacteriia) and cvE6 (Verrucomicrobiae), along with sequences assigned to the families Sandaracinaceae (Myxococcia) and Micropepsaceae (Acidobacteriia) (Fig. 4a). The genera *Sphingomonas*, *Caulobacter*, *Rudea*, *Rhodanobacter* and further taxa belonging to the families Rhodanobacteraceae, Fimbriimonadaceae and Elsteraceae, were significantly increased across the LCAP as well as cellulose and PHBV groups.

**Figure 4.**
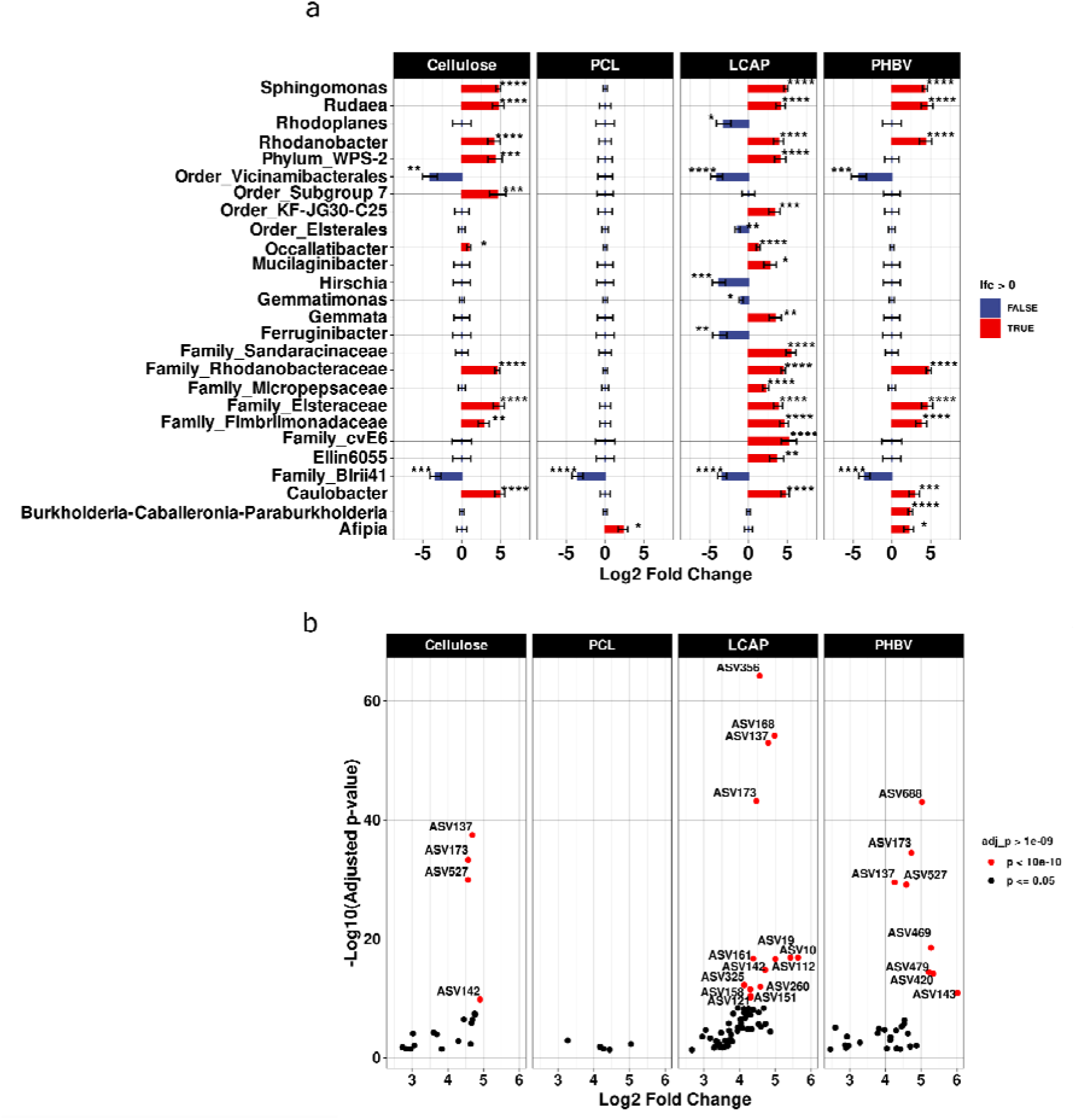
Log-fold change (LFC) analysis for differentially abundant bacterial taxa at the genus level (a) and ASV level (b) in the cellulose and PCL-, PHBV- and LCAP-bioplastic degrading microcosms. Non-spiked microcosms (blanks) and the inactive HDPE-spiked microcosms were grouped together as reference. Likewise, all microcosms treated with the three types of LCAPs were grouped together, while replicates of the cellulose-, PCL- and PHBV-treatments formed separate groups. To reduce the impact of sequencing artifacts and extremely rare ASVs, only ASVs with a count of ≥100 across 10% of samples were included in the analysis. Red bars (a) or dots (b) indicate a significant increase in a taxon’s abundance compared to the control condition. Blue bars indicate a reduction (a) and black dots (b) indicate taxa with an insufficient P-value. Taxa are displayed at the genus level (a) if classification was possible with sufficient confidence, otherwise the highest reliable taxonomy level is indicated. Statistical significance of the LFC is indicated next to the bars (*: P ≤ 0.05, **: P ≤ 0.01, ***: P ≤ 0.001, ****: P ≤ 0.0001).

At the ASV level and for the LCAP group, the most significant increases (p < 10e^-10^) were observed for an ASV belonging to the order Acidobacteriales (ASV 356) and an ASV classified as *Pajaroellobacter* within the order Polyangiales (ASV 168), which were exclusively observed for the LCAP group. Additional ASVs with highly significant increases, observed exclusively for the LCAP group, belonged to the Micropepsaceae (ASVs 10,19,112,158), *Terracidiphilus* (ASVs 161&325), *Oaccalatibacter* (ASV 151) and Elsteraceae (ASV 260). Highly significant increases were also observed for ASVs of *Sphingomonas* (ASV 137), Rhodobacteriaceae (ASV173) and *Caulobacter* (ASV142), however, those were also significantly increased in the PHBV and/or the cellulose groups. ASVs that increased only in the PHBV group were exclusively related to the BCP group (ASVs 688,479,469,420,143), while an ASV related to Tepidispherales (ASV 527) was also increased in the cellulose group. No uniquely enriched ASVs were observed for the cellulose group, while the PCL group showed no ASVs with significance levels p < 10e^-10^.

### Metagenomics reveals candidate genes for putative extracellular LCAP depolymerase enzymes

To reveal genes for potential LCAP-depolymerizing enzymes in the microcosms, the DNA samples from the microcosm replicates were pooled and used for metagenome shotgun sequencing in comparison to the pooled DNA samples of the PCL, PHBV, HDPE and cellulose microcosms; the non-spiked controls were not analyzed further.

Analysis of the resulting metagenomic contigs identified 2,624 unique candidate enzyme sequences with homology to known plastic-degrading enzymes, based on Hidden Markov model (HMM) motifs of enzyme sequences stored in the PlasticDB.^17^

The dataset was further analyzed via Principal Component Analysis (PCA), based on the normalized coverage of the contig encoding the respective protein, resulting in a clear clustering according to the polymer treatment with LCAP samples grouping closely together, while all other treatments occupy distinct coordinates, suggesting a marked difference in coverage of contigs encoding candidate plastic depolymerase enzymes (Fig. 5a). A PCA visualization displaying the individual proteins revealed that out of the 2,624 identified candidate enzymes, only a subset exhibited strongly differing contig coverage, highlighting their treatment-specific enrichment (Fig. 5b). All candidates with a negative value on the PC2 axis and a value >5 on the PC1 axis, associated with increased coverage for the LCAP samples, were manually curated and selected for further analysis based on three criteria: (*i*) HMMer search results with an e-value of <10e^-5^ and (*ii*) a BlastP e-value of <10e^-10^, and (*iii*) possession of a signal peptide for secretion of the candidate enzyme.

**Figure 5.**
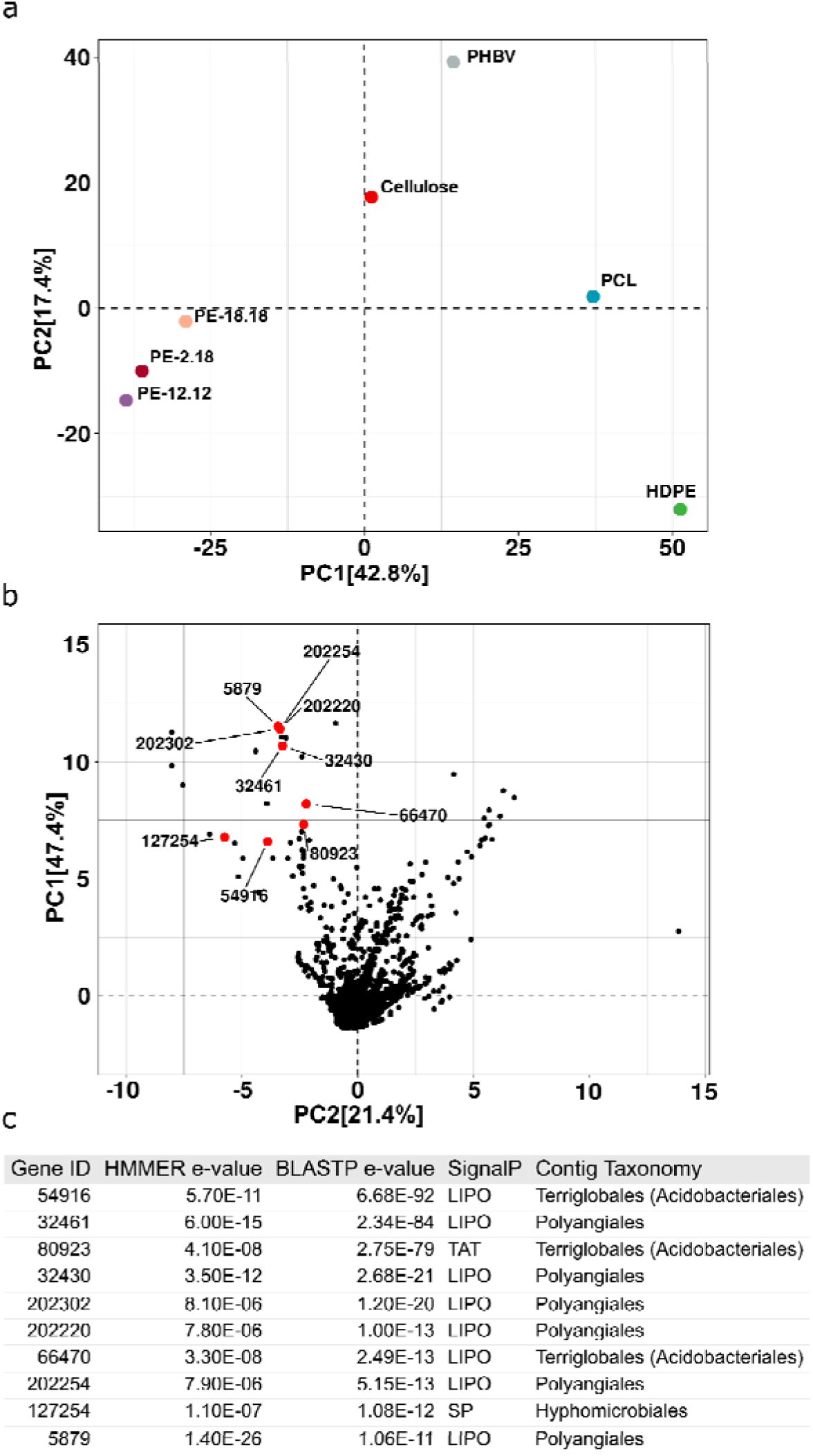
Principal Component Analysis (PCA). Shown are PCA results based on the normalized coverage of the contig encoding the putative depolymerase proteins, visualized **(a)** by microcosm treatment and **(b)** as volcano plot, and illustrating the clustering of individual proteins with the ten highest scoring matches highlighted and labelled with their respective gene identifiers (Gene ID number). Percentages given at the axes indicate the dissimilarity explained by the respective axis. **(c)** Gene ID, HMMer and protein Blast search e-values, along with predicted signal peptide and contig taxonomy of the top-matches highlighted in the volcano plot are shown in a tabular form.

The taxonomic affiliation of the top-10 best matches was estimated based on contig taxonomy, indicating that four were affiliated with the order Terriglobales (Acidobacteriales in the Silva taxonomy), five with the order Polyangiales and one with the order Hyphomicrobiales (Fig. 5c). Out of these matches, four exhibited BLASTP e-values below 1e^-50^, suggesting a high sequence homology to previously described polymer-degrading enzymes in the PlasticDB database, while one was found to not correspond to a complete protein and was excluded from further analysis. This left three candidate enzymes, GeneIDs (GID) 32461, 54916 and 80923, as the top-most striking candidates, since their taxonomic affiliation with the orders Acidobacteriales and Polyangiaceae closely matched the two most significantly increased ASVs in the long-chain PE group, ASV356 and ASV168, as indicated by the results of the LFC analysis of the 16S rRNA-gene sequencing data (Fig. 4b).

Candidate GID32461 showed the highest sequence similarity (36.5%, E-value: 5.4e^-75^) to the *p*-nitrobenzylesterase of *B. subtilis* (PlasticDB-ID 134), reported to hydrolyse PET.^35^ The closest relative in Uniprot was the carboxylic ester hydrolase of *Sorangium cellulosum* strain So ce56 (Uniprot ID A9FEC8).^36^ The search of the Interpro database grouped GID 32461 into the carboxylesterase family (IPR002018) within the α/β-hydrolase fold superfamily (IPR029058), further suggesting a potential esterase activity of GID32461.

Candidate GID80923 also shared high sequence homology with PlasticDB entry 134, as well as with entry 79, with 38.8% and 34.2% sequence identities and e-values of 2.7e^-72^ and 2.1e^-72^ respectively. A Uniprot search revealed a high sequence identity of 75% to a carboxylic ester hydrolase of *Alloacidobacterium dinghuense* with entry ID A0A7G8BC64. As with GID32461, GID80923 can be assigned to the carboxylesterase family (IPR002018).

Interestingly, the closest match to candidate GID54916 in Uniprot was a β-lactamase enzyme of *Granulicella* sp. WH15 with Uniprot ID A0A7L5A8I1,^37^ while a metagenome-sourced esterase, reported to depolymerize PLA,^22^ was the closest match in the PlasticDB database, with 42% sequence identity and an E-value of 1.3e^-92^ (PlasticDB entry 158). Both the GID54916 candidate and the PlaticDB entry 158 possess a β-lactamase/transpeptidase related domain (IPR012338).

All three LCAP-depolymerase candidates possess signal peptides for secretion, as predicted by SignalP-6.0.^38^ Presence of a secretion signal was considered a prerequisite for putative LCAP-degrading enzymes, due to the necessity of extracellular depolymerization. GID80923 is predicted to be secreted via the twin-arginine translocation (TAT) pathway, while GID32461 and GID54916 are predicted to be secreted via the general secretion (Sec) pathway. Additionally, GID32461 and GID54916 were found to contain a lipobox motif, suggesting that these proteins are lipidated following signal peptide cleavage, and subsequently anchored to the cell membrane, rather than being secreted freely into the extracellular or periplasmatic space (see also Discussion).^39^

To confirm the validity of the conducted analysis, we decided to further investigate the LCAP depolymerase potential for one of the top-three candidate enzymes, GID54916, both computationally and through heterologous expression and activity testing, as reported in the following.

### Modelling of the protein structure and substrate docking simulations for the predicted LCAP-depolymerase candidate GID54916

The AlphaFold 3-predicted structure of GID54916 revealed strong similarity to the esterase EstB from *Burkholderia gladioli*, belonging to the family VIII esterases, which share several conserved features with class C β lactamases (Fig. 6).^40^ Family VIII esterases exhibit the esterase atypical catalytic S-X-X-K motif, which matches the catalytic motif of class C β-lactamases.^20,40^ Other features shared between esterase-family VIII enzymes and class C β-lactamases are the S-D-N loop and the K-T-G box that contribute to the shaping of the catalytic cavity.^20,40^ While the S-X-X-K motif harbors the nucleophilic serine and the catalytic lysine, the Y of the S-D-N loop is the third residue of the catalytic triad. The K-T-G box of class C β-lactamases corresponds to the W-X-G motif of family VIII esterases (H-X-X for subfamily 3).^40^ A multiple sequence alignment of GID54916 and its closest PlasticDB matches, together with EstB and additional family VIII esterases, as well as class C β-lactamases, confirmed that GID54916 has all the above mentioned conserved regions, thereby grouping it within the type VIII esterase family (Fig. S5).

**Figure 6.**
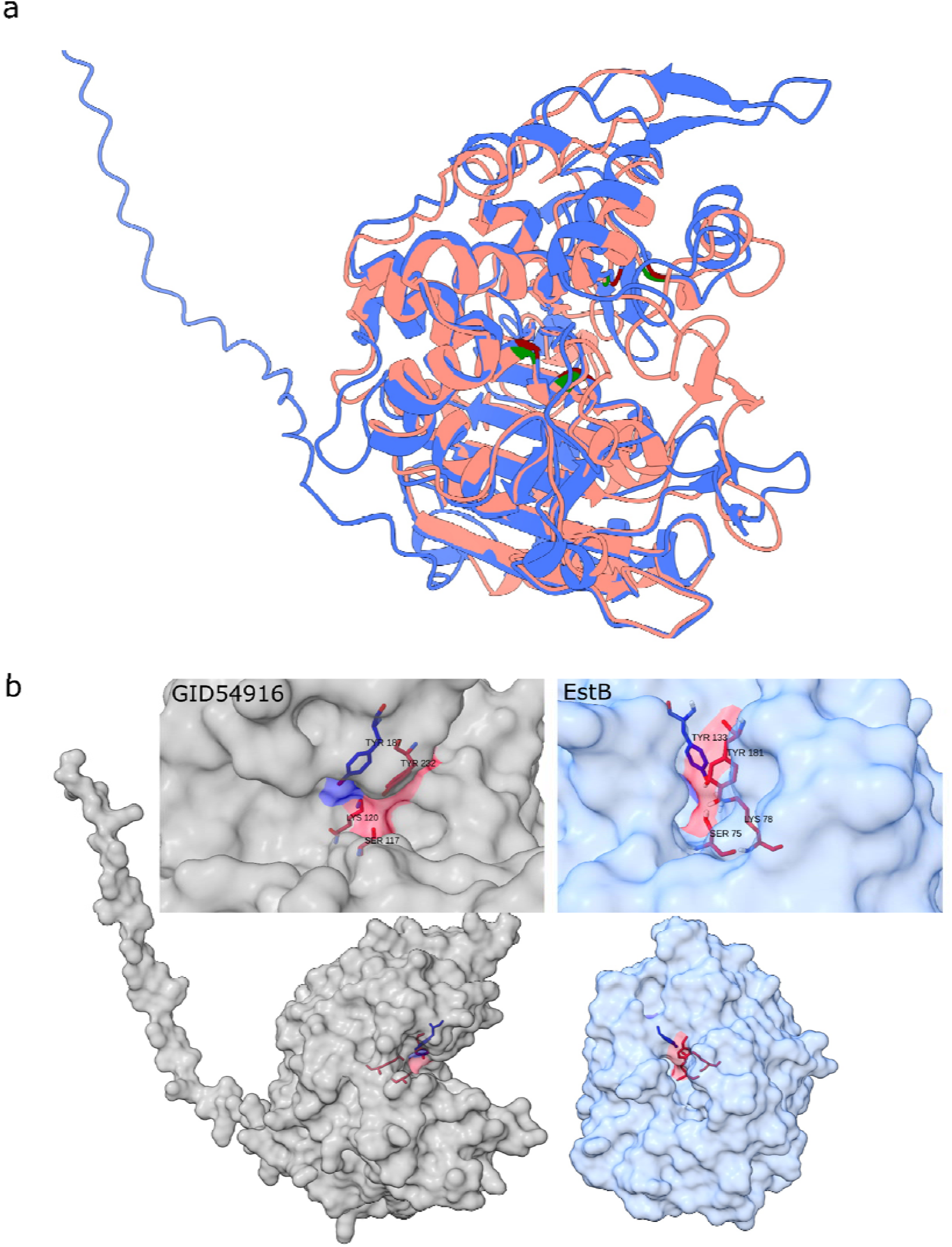
Predicted protein structures (a) and surface models (b) of GID54916 in comparison to its close homologue, characterized esterase EstB of *Burkholderia gladioli*. (**a**) Overlay of the two protein structures, with GID54916 in blue and EstB in pink. The conserved residues forming the active sites are marked in green for EstB and red for GID54916. (**b**) Comparison of the surface structures of GID54916 (light-grey) and of EstB (light-blue), illustrating a much more widely surface-exposed active site in a large accessible binding-groove as predicted for GID54916, rather than a tunnel as for EstB. The active-site surfaces are highlighted in red.

While family VIII esterases are known for their broad substrate spectrum and structural similarity to β lactamases, these enzymes typically lack the ability to hydrolyze β-lactams,^40^ as is the case for EstB, due to the spatial limitations in its active site, which is constrained by a tunnel-like structure that cannot accommodate the comparatively bulky β-lactam molecules.^20^ Interestingly, for the structural model of candidate GID59416, the active site was significantly more exposed, forming a large accessible binding-groove and resulting in a protein with a striking ‘pac-man’-like structure, rather than a tunnel (Fig. 6b and 7a).

**Figure 7.**
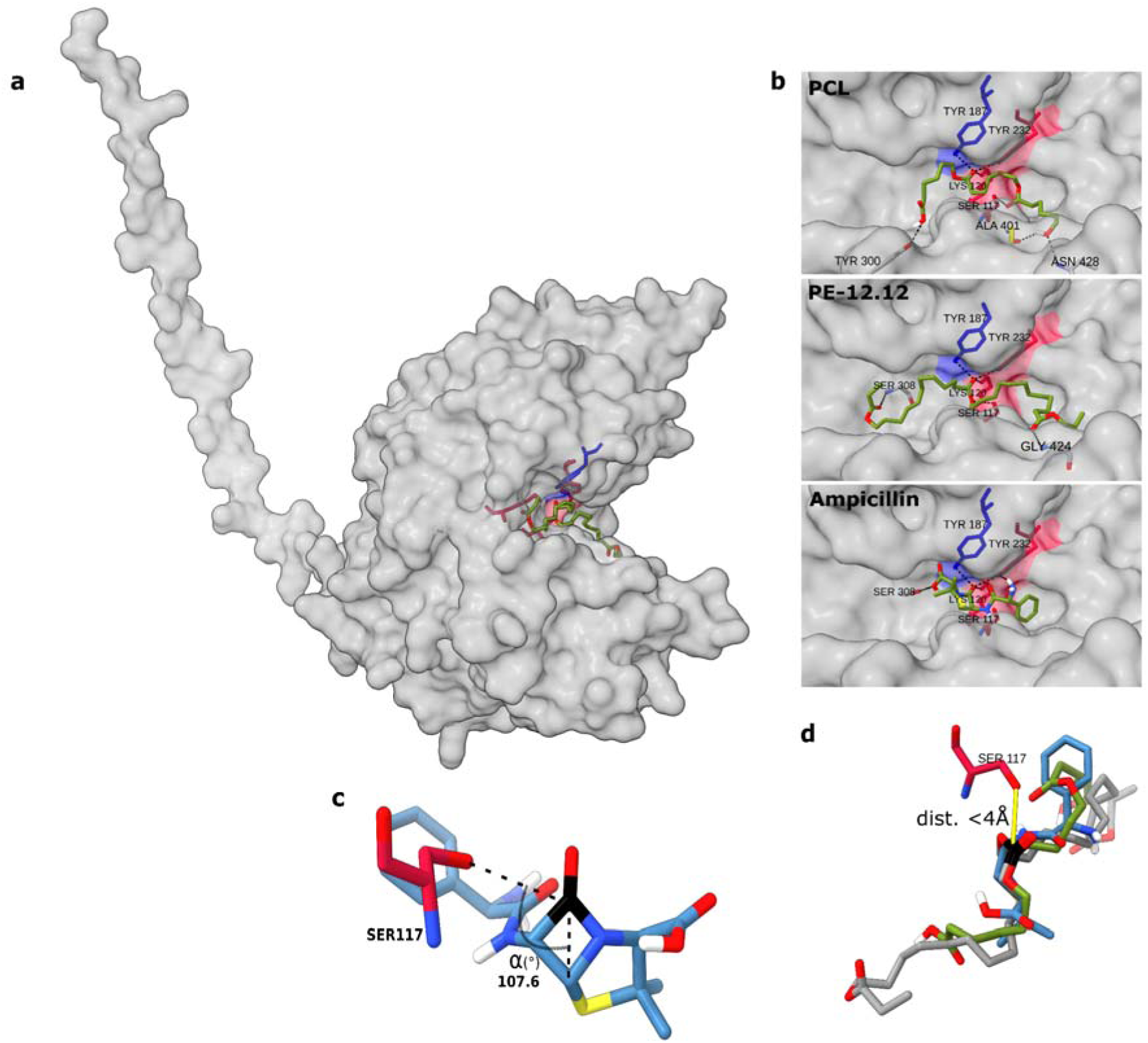
**Alphafold-predicted surface structure of GID54916 with bound ligands**. The catalytic triad residues (Ser117, Lys120 and Tyr232) are highlighted in red and Tyr187 in blue. (**a**) The PE- 12,12 dimer bound to the enzyme surface is depicted in green with red sticks indicating the oxygen atoms of ester bonds. (**b**) Close-up of PCL, PE-12,12 and Ampicillin bound to the GID54916 active site. Potential hydrogen bonds formed between the ligands and catalytic enzyme residues are indicated by black dotted lines. (**c**) Visualization of the attack angle of Ser117 towards the carbonyl carbon of the Ampicillin β-lactam ring. (**d**) Overlay of the binding conformations of PE-12,12, Ampicillin and PCL with the carbonyl carbon of the esters (PE-12,12, PCL) or β-lactam group highlighted in black. The yellow line indicates the distance of the carbonyl carbon to the nucleophilic Ser117.

Docking simulations using the polyester ligands PE-12,12 (dimer) and PCL (trimer), as well as the β-lactam antibiotic Ampicillin, resulted in docking solutions with similar binding energies, that placed the respective ligand in close proximity to the active site (Fig. 7). The observed docking solutions furthermore resulted in a favorable geometry for catalytic activity for all ligands. The carbonyl carbon of the ester or β-lactam group are in close proximity to the nucleophilic Ser117 (<4 Å), while the oxygen atom of the functional group is in close enough proximity to Lys120, Tyr232 and Tyr187 to form hydrogen bonds, enabling the activation of the carbonyl carbon and the stabilization of the oxyanion during the catalytic process (Fig. 7b and d). Additionally, the attack angle for the β-lactam carbonyl likewise appeared to be ideal for nucleophilic attack, closely matching the canonical Bürgi-Dunitz angle of 107° (Fig. 7c).^41^ Hence, the structural modeling and docking simulations suggested that GID54916 likely possesses a broad substrate spectrum thanks to a widely accessible active site that may be able to accommodate and hydrolyze large substrates, such as β-lactam antibiotics and polyesters.

### Heterologous expression of GID54916 and enzyme activity testing

To confirm the *in-silico* predicted depolymerization of LCAP and hydrolysis of β-lactams by candidate GID54916, a soluble variant of the protein (without secretion signal and lipobox motive) was heterologously expressed, purified and the quality of the preparations examined by SDS-PAGE (Fig. S6). Esterase activity of the purified protein was confirmed in a colorimetric reaction using p-nitrophenyl butyrate as the substrate.

Subsequent enzyme assays with the LCAP PE-2,18 as substrate (powder) demonstrated octadecanedioate monomer release over time, while reactions without enzyme and with heat-denatured enzyme showed no octadecanedioate formation (Fig. 8a). The highest production was observed within the first 4 h, for which a specific activity of 0.5 U mg^-1^ was calculated. The other monomer of PE-2,18, ethylene glycol, was also released in comparable proportions, as followed by end-point determination in parallel reactions (Fig. 8b), with 21.8 ± 3.1 µM octadecanedioate and 18 µM ethylene glycol after 24 h and 18 h, respectively.

**Figure 8.**
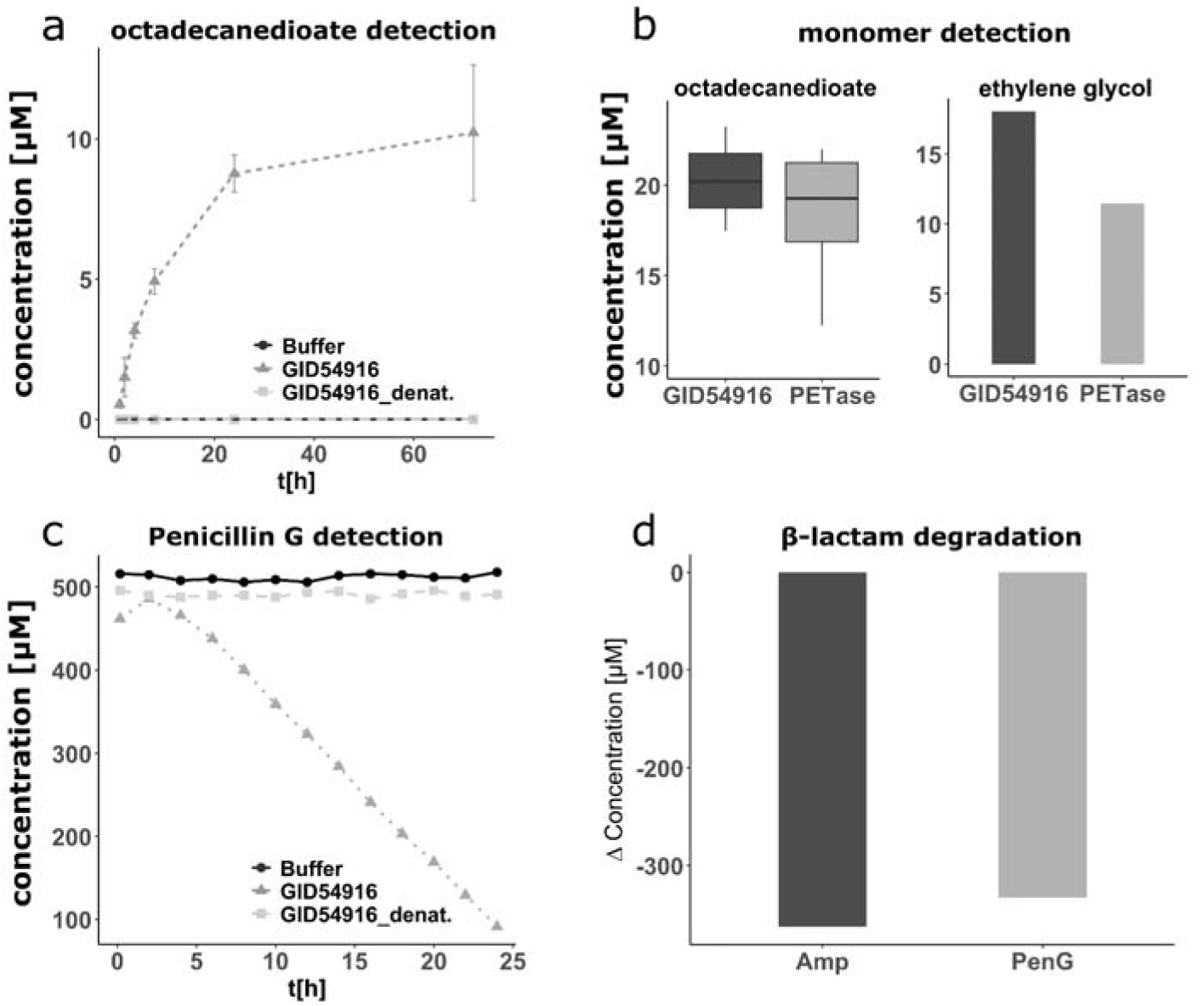
LCAP depolymerization and hydrolysis of β-lactam antibiotics by recombinantly produced GID54916 enzyme. **(a)** Time course of octadecanedioate monomer production in incubations of GID54916 with a powder of the LCAP PE-2,18 as substrate over 72 h, as compared to heat-denatured GID54916 and a buffer control. Reactions contained 5 mg of PE-2.18 powder and 2.5 µM of enzyme. The formation of octadecanedioate in the supernatant was analyzed by HPLC-MS in triplicate incubations; standard deviation is indicated by error bars. **(b)** End-point concentrations of both PE-2,18 monomers, octadecanedioate and ethylene glycol, after incubation of PE-2.18 with GID54916, or with recombinantly produced *Ideonella sakaiensis* PETase for comparison. Each 5 mg PE-2,18 and 2.5 µM enzyme were used. Octadecanedioate in the reactions (n = 3) was determined by HPLC-MS and ethylene glycol by HPLC-RID in a parallel reaction (n = 1), after 24 h and 18 h, respectively. **(c)** Time course of Penicillin G hydrolysis by GID54916 over 24 h, as determined by HPLC-DAD. The reaction contained 500 µM Penicillin G and 2.5 µM GID54916 and was analyzed every 2 h for 24 h. Control incubations with heat-denatured GID54916 or only buffer were included. **(d)** Concentration decreases of β-lactam antibiotics Penicillin G or Ampicillin after incubations with GID54916 for 24 h. Reactions contained 2.5 µM GID54916 and 500 µM Penicillin G or Ampicillin, and the antibiotics concentrations were determined before and after the incubations.

Reactions with recombinant *Ideonella sakaiensis* PETase and PE-2,18 produced similar amounts of monomers (Fig. 8b), suggesting comparable activities of the two depolymerases under the experimental conditions used. GID54916 also depolymerized PCL, however at a significantly higher rate (Fig. S7; up to approx. 2 mM caprolactone in 24 h), likely owing to a higher frequency of ester bonds and a low molecular weight and crystallinity of PCL compared to PE-2,18.

Additional tests confirmed β-lactam hydrolysis by GID54916, demonstrating Penicillin G disappearance in incubations with active GID54916, whereas incubations with heat-denatured GID54916 or without enzyme showed no activity (Fig. 8c & S9). The Penicillin G disappeared nearly completely within 24 h, for which a specific activity of 8.92 U mg^-1^ was calculated (Fig 8c, Fig. S10). A second β-lactam antibiotic, Ampicillin, was also hydrolyzed, as shown by end-point determination in parallel reactions (Fig. 8d). The inactivation of Ampicillin was furthermore confirmed by disc diffusion assays, for which GID54916 was added to Ampicillin-containing filter discs on agar plates, resulting in a clear inactivation of the inhibitory effect of Ampicillin on *E. coli* K12 (Fig. S11).

## Discussion

This study was initiated to investigate the microbial degradation of different long-chain aliphatic polyester (LCAP) plastics materials (PE-18,18, PE-12,12 and PE-2,18) under mesophilic conditions in forest soil, and lead to the discovery of GID54916 (termed LCPH1) as a novel, most likely membrane-associated family VIII esterase that catalyzes long-chain aliphatic polyester (LCAP) depolymerization, as well as β-lactam hydrolysis, in a rare dual functionality.

Microcosm experiments demonstrated the near-complete mineralization of the LCAPs PE-2,18 and PE-18,18 within 8–10 months, at rates similar to those observed for the commercial biodegradable plastics PHBV and PCL. In contrast, PE-12,12 degradation reached a plateau at ∼20% of polymer-carbon conversion to CO_2_, despite comparable crystallinity across the different polymers. Also the chain length (molecular weight) of the different LCAPs (PE-18,18 > PE-2,18 > PE-12,12) may not explain the limited degradation of PE-12,12. Degradation of LCAPs was previously demonstrated under industrial, thermophilic composting conditions and in an agricultural soil under mesophilic conditions.^32,33^ The latter study could demonstrate significant degradation only for shorter-chain LCAP variants (while they exhibit inferior material properties), whereas only a low extent of degradation of PE-2,18(±2) was observed.^33^ Certainly, the different degradation kinetics observed across different degradation studies, using different sources of microbes and incubation conditions, and different LCPE preparations, warrants further investigation.

Furthermore, our experiments highlighted that limitations in essential nutrients can restrict plastic biodegradation rates. Supplementing N, P, S and trace metals to the forest soil improved degradation, suggesting that natural environments, which are often the recipient of post-consumer plastics, might not exhibit their full degradation capacity due to nutrient co-deficiencies. This is supported by studies showing the positive effect of nitrogen-fixing bacteria on polymer degradation, as reported for poly(butylene succinate-co-adipate) (PBSA),^42^ as well as improved degradation of polymers when additional N was supplied in the polymer matrix or through the inclusion of natural N-containing fillers.^43,44^ However, the most profound hindrance of efficient microbial degradation of bioplastics, was deemed to be insufficient water availability, as concluded from our field trial in forest soil during a very dry year (2022 in Germany).

The amplicon and metagenomic sequencing of the microcosm experiments identified taxa and candidate enzymes likely involved in LCAP degradation. Taxa related to Acidobacteriales and Polyangiales, both abundant in soil ecosystems, were significantly enriched in LCAP-treated samples, suggesting specialized roles in polymer degradation. Across samples, 2,624 coding genes were identified with significant similarity to HMM motifs obtained from reported plastic depolymerases (hydrolases). PCA based on normalized contig coverage enabled prioritization of candidate genes enriched in LCAP-treated samples. Nine of the top ten candidates showed contig-level affiliation with Acidobacteriales or Polyangiales, consistent with amplicon-based community profiles.

Among these, GID54916 (Acidobacteriales), encoding LCPH1, was selected for further analysis due to its high sequence similarity to a previously reported depolymerase (PlasticDB entry 158) and its prediction as a secreted lipoprotein with structural similarity to type C β-lactamases. In Gram-negative bacteria, lipoproteins are typically anchored to the inner leaflet of the outer membrane,^45^ although recently improved understanding of Gram-negative lipoproteins suggests that both cell surface exposure, i.e. anchoring to the outer membrane leaflet, and further secretion of periplasm-facing lipoproteins via outer membrane vesicles (OMVs), may be more prevalent than previously estimated.^46–51^

This finding piqued our interest since membrane association of known plastic-depolymerizing enzymes appears exceptionally rare. To the best of our knowledge, the only reported lipoprotein involved in plastic degradation is the *I. sakaiensis* MHETase, which degrades the water-soluble PET dimer MHET (mono-2-hydroxyethyl terephthalic acid).^52^ While there appear to be no reports of extracellular plastic-depolymerizing lipoproteins to date, these attributes might have been overlooked or considered irrelevant. For instance, the membrane-associated polyurethane esterase (PudA; PlasticDB entry 13) contains a lipobox motif but was not reported as a lipoprotein.^53^ Apart from lipoproteins, other membrane associated plastic-degrading proteins are likewise scarcely reported, such as the PET-degrading transmembrane esterase in Gram-positive *Rhodococcus pyridinivorans* P23.^15^ Furthermore, the involvement of OMVs in the degradation of polyurethane oligomers was demonstrated, which implies another membrane-associated secretion mechanism involved in plastic degradation.^16^ These observations suggest that depolymerases associated to the outer cell surface (ectoenzymes) may be a yet underrecognized strategy in plastic-degrading natural microbial communities, while this strategy is widespread among plant polymer-degrading bacteria, which frequently employ cellulosomes associated to the cell surface,^54^ as well as OMVs for plant cell wall degradation.^55–57^ Considering that depolymerizing enzymes for natural and synthetic polymers often belong to the same enzyme families, the adaptation of a similar process for synthetic polymer degradation appears likely. Such a functional display of the depolymerases on the cell surface would (*i*) reduce loss of the enzymes to the extracellular space (especially in aqueous environments), (*ii*) result in an improved efficiency due to high local concentration of enzymes at the bacteria–plastic interface, and (*iii*) ensure catalysis close to the cell surface and thus efficient monomer uptake, limiting potential loss of the monomers to cheaters, i.e., to commensal, monomer but non-polymer degrading microbial community members that are not investing in the synthesis of extracellular depolymerases.

Indeed, such a catalytic advantage was experimentally demonstrated, showing that a PETase functionally displayed on the surface of yeast cells and combined with hydrophobin- mediated cellular plastic adhesion, resulted in an up to 329-fold increased efficiency compared to freely dissolved PETase.^58^

Such a functional display of the depolymerases on the cell surface, and/or close to the cell surface (OMVs), may also have resulted in the single-cell-shaped voids (‘sinkholes’) in the plastic films as retrieved from the forest soil after our field trial, and as observed by electron microscopy (Fig. 1 & Fig. S2). Hence, this observation may represent a clear visual evidence of a direct catalytic activity of the microbial cell surface at the cell-plastic interface.

Notably, several further genetic features, in addition to the signal sequence and lipbox motif of LCPH1, support an outer-membrane association and potential involvement of OMVs in its secretion, while a direct experimental confirmation of LCPH1’s cellular localization must await future experiments: The genomic context of the contig encoding LCPH1 includes genes for LolC/LolE-like permease components that traffic lipoproteins to the outer membrane, a cell- wall-remodeling enzyme MepM/NlpD (murein endopeptidase or amidase activator) implicated in envelope dynamics and OMV biogenesis,^59^ and a GGDEF-domain protein, indicating involvement of c-di-GMP signaling, a key regulator of biofilm formation, surface attachment, and coordinated secretion responses, including OMV formation.^60,61^ Additionally, directly adjacent to the LCPH1 gene, the contig encodes a TonB-dependent receptor for potential uptake of complex degradation products in a conformation previously suggested for nutrient acquisition in conjunction with surface-exposed substrate-binding lipoproteins in *Bacteroides fragilis*.^46,62^ TonB-dependent receptors were also found to be enriched in in OMVs.^56,63^ Together, the contig-level co-occurrence of these features, along with carbohydrate-active enzymes in the same region, support a model in which LCPH1 is surface-exposed and may be secreted via OMVs, along with synergistic enzymes, during colonization of plant material or other polymeric substrates like plastics, contributing to extracellular degradation and efficient substrate utilization.

In addition to the interesting microbial-ecological and genomic contexts, the predicted structural features of LCPH1 highlight intriguing functional aspects. Structural analysis of LCPH1 revealed striking similarities to family VIII esterases, and particularly EstB from *Burkholderia gladioli*, leading to our assignment of LCPH1 to this enzyme family (Fig. S5). Family VIII esterases are known for their broad substrate range and are characterized by their similarity to type C β-lactamases.^40^ Despite these similarities, β-lactams are not typically within the substrate spectrum of family VIII esterases, for instance due to steric hindrance in the active site, as was demonstrated in case of EstB.^20^ A structural comparison of LCPH1 to EstB, however, revealed an intriguingly wide, surface-exposed active site of the LCPH1 protein in a ‘pacman-like’ shape, with the potential to bind large substrates (Figs. 6b and 7a).

This was confirmed *via* substrate docking simulations, which showed the favorable binding of polyester chains of PCL and PE-12,12, as well as the β-lactam Ampicillin in the active site of GID54916 (Fig. 7).

These computational predictions were confirmed experimentally, demonstrating both the LCAP depolymerase and β-lactamase activity of the enzyme (Fig. 8). While reports of β-lactamase activity among family VIII esterases keep increasing,^24–30^ this is the first instance where for a family VIII esterase both, plastic depolymerase and β-lactamase activity, is reported.^21–23^ However, this overlap in substrate spectrum was also recently reported for a metagenome-derived esterase PET40, which exhibited both PET depolymerase and β-lactamase activity, despite lacking the canonical β-lactamase fold.^19^

This dual activity of LCPH1, and of PET40, poses interesting questions regarding the ecological roles of these enzymes and potential unexpected consequences of plastic litter in the environment. Based on our results, it is likely that LCPH1 provides resistance to penicillin-like β-lactams, while also conveying additional fitness by enabling the utilization of (synthetic) polyester substrates. This duality might therefore result in a selection of antibiotic-resistant microbiota as a result of plastic exposure. Interestingly, studies have frequently reported the increased occurrence of antibiotic resistance genes on plastic debris, including β-lactamase marker genes, which was attributed mostly to plastics being a favorable surface for biofilm formation and horizontal transfer of ARGs, as well as sorbents for antibiotics and other contaminants.^9–13^ With regard to the current results, it is tempting to speculate on an additional factor: the potential overlap in the substrate range between certain secreted and surface-presented plastic depolymerases and antibiotics-degrading enzymes.

In summary, our study demonstrates the mesophilic degradation of LCAP bioplastics by native forest soil microbiota, while identifying nutrient and water availability as key environmental constraints of post-consumer plastic receiving environments. The subsequent sequencing analysis resulted in the identification of LCPH1, the first experimentally confirmed family VIII esterase with bifunctional polyesterase and β-lactamase activities. This rarely reported overlap in the substrate spectrum may contribute to the frequently observed enrichment of ARGs in the plastisphere, while the predicted surface-presented and OMV-mediated secretion of LCPH1 introduces a new microbial-ecological perspective on plastic degradation.

## Methods

### Polymer materials

Poly(3-hydroxybutyrate-co-3-hydroxyvalerate) (PHBV) was purchased from Natureplast (grade: PHI002, Mondeville, France). Polycaprolacton (PCL) was purchased from Sigma-Aldrich (Product ID: 440752, St. Louis, MO, United States). Cellulose powder (Sigmacell Type 20) was likewise purchased from Sigma-Aldrich. Long chain aliphatic polyester (LCAP) bioplastics PE-2,18, PE-12,12, and PE-18,18 were synthesized in house as previously reported.^64^ The numbering indicates the chain length, that is number of carbon atoms in the diol and dicarboxylate derived monomeric units. High density polyethylene (HDPE) was likewise synthesized in house to avoid potential contamination with industry-typical additives, using ethylene, 5 mL of toluene as solvent and 5 µmol of naphtyl-MeOBIPh-Ph as a catalyst. For the degradation test in forest soil *in situ*, plastic films of PE-18,18 and PE-12,12 material were used, as produced previously described.^64^ For the degradation test with forest soil incubated in laboratory microcosms, plastic materials PHBV, HDPE, PE-2,18, PE-12,12, and PE-18,18 were used, each milled to a fine powder after immersion of samples in liquid nitrogen using a Retsch ZM 200 Ultra Centrifugal mill (Retsch GmbH, Haan, Germany) at 14,000 rpm, equipped with a ring sieve with a mesh size of 1 mm.

### *In situ* degradation assay

Film strips of approx. 1 cm x 3 cm in size were mounted in between plastic grids in frames.^65^ For deployment of the materials into forest soil at the Botanical Garden of the University of Konstanz (mixed forest, predominantly beech and pine; coordinates, 47.691552, 9.179949), the top layer of litter was carefully removed from the soil, as well as the layer of partly decaying organic material (duff) up to a depth of approx. 10 cm into the humus layer; smaller plant roots had to be removed. The frames were placed onto the humus material and then covered approximately 2 cm deep with sieved (mesh size, 5 mm) humus material as freshly taken from a closely located site. Finally, the duff material and the litter were placed back onto the soil. The frames were kept in the soil for over one year (2022) covering spring, summer, winter and following spring. After recovery, the films were fully dried at room temperature and examined by scanning electron microscopy.

### Scanning electron microscopy

LCAP films from the forest soil incubation were cut into 5 x 5 mm pieces with a scalpel and adhered to silver tape, each mounted onto carbon pads on aluminum stubs (12 mm diameter). These samples were then coated with a 6.5 nm layer of platinum using a sputter coater (Quorum Q150 R ES, Quorum Technologies, Lewes, United Kingdom). The morphological analysis of the film was conducted using a scanning electron microscope (Zeiss Auriga Cross Beam, Zeiss, Oberkochen, Germany), operated at 5 kV with the secondary electron detector.

### Microcosm degradation assay

Biodegradation of polymers in forest soil was assessed under laboratory conditions through determination of CO_2_-production as previously described.^34,43^ Briefly, 1g of forest soil was weighed into 25-mL culture vials and spiked with 50 mg of cellulose (control) or PHBV, PCL, HDPE, PE-2,18, PE-12,12, or PE-18,18. The polymer-spiked soil samples were prepared in triplicate and mixed thoroughly before the addition of sterile ddH_2_O to achieve a moisture content equivalent to 70% of the soil’s water holding capacity (WHC). As a reference for the basal respiration in the forest soil, non-polymer spiked soil samples were prepared, likewise in triplicate. The vials were closed with modified butyl-stoppers equipped with CO_2_-traps filled with 0.1 M NaOH. The sample vials were incubated at 30 °C and the NaOH sampled regularly. The CO_2_ captured in the NaOH-solution was quantified in a TOC-L (Shimadzu, Kyoto, Japan) and the results used to generate cumulative degradation curves. The concentration of the NaOH-solution was increased to 0.5 M as saturation was observed at early incubation time points. To counteract suspected nutrient limitation during degradation, 200 µL of a nutrient solution were supplied after 66 days. Nutrient addition provided 2 mM potassium phosphate, 0.125 mM magnesium sulfate, 10 mM ammonium chloride and µM-concentrations of various trace metals (see Table S1), as calculated based on the volume of total water content of the soil microcosms. Through nutrient addition, the moisture content was increased to 80% of the soil WHC.

### DNA extraction and PCR

DNA was extracted from all samples after 164 days of incubation using a cetyltrimethylammonium bromide (CTAB)-based method adapted from Larsen and collegues.^66^ To avoid PCR inhibition by soil constituents such as humic acids, the DNA was additionally cleaned using polyvinylpolypyrrolidone spin-columns before down-stream applications.^67^ The 16S rRNA gene was amplified using primers Bakt341F (CCTACGGGNGGCWGCAG) and Bakt805R (GACTACHVGGGTATCTAATCC), targeting the V3 and V4 region of the gene.^68^ The internal spacer region 2 (ITS2) was amplified using primers fITS7 (GTGARTCATCGAATCTTTG) and ITS4 (TCCTCCGCTTATTGATATGC).^69^ All primers were equipped with the appropriate adapters for Illumina sequencing platforms.

### Amplicon sequencing analysis

The 16S rRNA amplicons were further processed as per the Illumina “16S Metagenomic Sequencing Library Preparation” protocol (Illumina, San Diego, CA, United States). Library preparation and sequencing on the Illumina MiSeq platform were performed by Eurofins genomics (Konstanz, Germany). The sequencing reads were processed in R (v.4.3.2) utilizing packages dada2 (v.1.26.0) as described in a previously reported analysis pipeline.^70^ Taxonomy was assigned to ASVs using the Silva database version 138.1 for bacterial 16S rRNA amplicons and the UNITE ITS database version 9 for fungal ITS amplicons.^71,72^ Microbial community analysis was performed in R using packages pyhloseq (v.1.42.0),^73^ Biostrings (v.2.66.0),^74^ ape (v.5.7.1),^75^ and vegan (v.1.8.8).^76^ Log-fold change analysis (lfc) was performed using R package ANCOMBC2 (v.2.0.2) with default settings and “struc_zero” set to FALSE.^77^ For lfc analysis, datasets from samples supplied with LCAP-materials were grouped together, while datasets of Cellulose, PCL, and PHBV treated samples were considered as independent groups. Datasets of samples without plastic addition were used as a reference group together with datasets of samples provided with HDPE, due to their similar community composition and lack of substrate-induced CO_2_ evolution in those samples. Prior to analysis, ASVs with less than 100 counts across 10% of samples were excluded to avoid over-estimation of rare taxa in the analysis. R packages ggplot(v.3.4.2) and ggsci(v.3.0.0) were utilized in data visualization.^78^

### Metagenome analysis

For metagenomic shotgun sequencing, the triplicate DNA samples of the respective treatments were combined and one sample per treatment was submitted for sequencing on an Illumina HiSeq device at Eurofins Genomics (Konstanz, Germany). Quality filtering of the resulting reads, metagenome assembly and analysis were conducted within the anvio (v.7.1) environment.^79,80^ Metagenome assembly was conducted using megahit (v. 1.2.9) with a minimum contig length of 1000bp. Mapping of the reads to the assembled contigs was done using bowtie2 (v.2.3.5).^81^ The anvio metagenome pipeline was further employed to create a contigs database, which includes the identification of open reading frames (ORFs) using Prodigal.^82^ Amino acid sequences for identified ORFs were exported and analyzed further.

The program “anvi-run-scg-taxonomy” was used to assign taxonomy to contigs based on single-copy core genes present in the metagenome with reference to the genome taxonomy database (GTDB).^83^ The program “anvi-profile” was used to obtain coverage information. The program ”anvi-estimate-scg-taxonomy” was used to estimate the taxonomic affiliation of bins based on single-copy core genes present in the assigned contigs.

### Identification of putative plastic degrading enzymes in the metagenome

Putative plastic degrading enzymes were identified by employing hidden markov models (HMMs) of previously reported plastic degrading enzymes, similar to the approach reported by Danso and colleagues.^84^ Amino acid sequences of all plastic degrading enzymes reported in the PlasticDB were downloaded and grouped by the plastics they were reported to degrade.^17^ Sequences of enzymes reported to degrade plastics polyethylene and polystyrene were combined. The grouped sequences were subsequently aligned using MAFFT (v.7.520),^85^ followed by the creation of HMM motifs using HMMer (v.3.1b2).^86^ Results were pre-filtered to include hits with e-values <10E-3. A local blastP (v. 2.12.0) search was conducted using the PlasticDB sequences as reference database to identify the closest database match.^87^ To obtain information about possible secretion motifs, pre-filtered sequences were analyzed using SignalP-6.^38^ Finally, replicate hits that matched multiple HMM motifs were removed, resulting in a dataset of unique metagenomic protein sequences with significant HMM-based similarity to previously reported plastic degrading enzymes. To obtain a more concise list of candidates potentially capable of LCAP depolymerization, the dataset was further analyzed based on normalized contig coverage to account for differences in total metagenome size between samples, in order to identify candidate enzymes with increased coverage in LCAP-treated samples. To this end, the sequences of candidate enzymes were analyzed via principal component analysis (PCA), using R packages FactoMineR (v. 2.11) and factoextra (v. 1.0.7),^88^ to identify the most differentially covered sequences in the dataset. Candidate sequences with a negative value on the PC2-axis and above a PC1-axis threshold of 5 were selected for manual curation based on the predicted presence of a secretion signal, a more stringent cut-off for the e-value of the HMMer search score (<10E-5), as well as their BlastP e-value (<10E-10). A list of candidate proteins is provided in the supporting information (excel file).

### Structural analysis and docking simulations

The structural model for LCPH1 used for docking simulations was obtained using the Alphafold3 web server.^89^ The active site of LCPH1 was predicted based on the obtained structural model using the ProFunc web server.^90^ Ligand structures used for modeling were downloaded from PubChem in the case of the β-lactam antibiotics Ampicillin and Penicillin. Structures of polyester oligomers (PE-12,12 dimer & PCL trimer) were obtained from their respective SMILES specification using Open Babel (v. 3.1.1).^91,92^ The protein and ligand structures were prepared for docking using Autodock Tools (wihtin MGLTools v. 1.5.7).^93^ The grid-box was placed centered on the active site serine and its size adjusted to fit the respective ligand. The maximum amount of active torsions was allowed for each ligand and docking was performed using the genetic Lamarckian algorithm. The search was conducted using a population size of 100, 2.5x10^5^ energy evaluations and a maximum number of 5x10^4^ generations in 100 individual runs.

In case of the oligomeric ligands, the docking simulation did not converge into clear clusters. Therefore, the best docking solution with close proximity of the central ester bond to the catalytic residues was chosen for refinement by repeated docking with reduced active torsions. For the PE-12,12 dimer, the number of active torsions was reduced from 30 to 16 and for the PCL trimer from 21 to 12, thereby reducing the complexity of the search. After refinement, clear clusters were obtained for the oligomeric ligands. The β-lactam ligands formed clear clusters without refinement being required.

For each docking simulation, the best scoring docking solution was exported for visualization and further analysis using ChimeraX (v. 1.8).^94^ Hydrogen bonds between ligand and protein, distances between the nuclephilic serine and the ligand carbonyl carbon, as well as the attack angle of the nucleophilic serine were determined using ChimeraX.

### Multiple sequence alignment

The LCPH1 amino acid sequence was aligned to homologous proteins from the PlasticDB database (IDs 91 and 158) as well as type C beta lactamases p99 of *Enterobacter cloacae* (PDB entry 1XX2) and BlaEC of *Escherichia coli* (PDB entry 9C6P), family VIII type esterases EstB of *Burkholderia gladioli* (PDB entry 1CI8), and metagenomic enzymes EH7 (PDB entry 7PP3) and Est-Y29 (PDB entry 4P6B). Multiple sequence alignment was conducted using the MAFFT web server using default settings except for the addition of further homologous sequences through the implementation of DASH.^95^ The sequences added by DASH for improved alignment were omitted in the final result. The alignment was visualized using Biopython and Matplotlib.^96,97^

### Heterologous expression of putative plastic degrading enzymes

The LCPH1-coding gene without its secretion signal, with C-terminal His-tag, and codon-optimized for heterologous expression in *E. coli* was ordered in a pET-26b(+) expression vector at BioCat (Heidelberg, Germany). The vector was transformed into *E. coli* expression strain Rosetta Gami 2 DE3, and the insert confirmed via restriction digest. For expression of the target enzyme, 50 mL over-night cultures were prepared using LB-medium supplemented with 50 mg/L Kanamycin (Kn_50_) and 20 mg/L Chloramphenicol (Cm_20_). Kn-resistance was conveyed by the pET-26b(+) vector, while Cm-resistance was conveyed by a Rosetta Gami 2 plasmid encoding rare t-RNAs. Over-night cultures were inoculated with transformed cells and grown at 37 °C and 200 rpm horizontal shaking in 200 mL Erlenmeyer flasks. The cultures were then used to inoculate 1 L cultures containing 950 mL of LB medium with additional 10 mM Glucose and Kn_50_ and Cm_20_. Glucose was added to prevent basal expression prior to induction by carbon catabolite repression.^98^ The cultures were then grown to an OD600 of 0.6-1.0 before induction with 0.4 mM IPTG. The cultures were then transferred to a cooled incubator for expression at 18 °C and 100 rpm for 18-20 h.

Cells were centrifuged at 5,000 x *g* for 15 min and washed twice with 50 mM PBS (pH 7.4, 300 mM NaCl). Cell pellets were frozen in liquid nitrogen and stored at -80 °C until further processing. For cell lysis the cells were resuspended in 50 mM PBS (pH 7.4, 300 mM NaCl) supplemented with 10 mM imidazole and 0.1% Triton-X. Triton-X was used to aid in protein solubility during the lysis and extraction process. Additionally, cOmplete™ ULTRA proteinase inhibitor tablets (Roche, Basel, Switzerland) and DNAse I were added to the lysis mixture.

Lysis was achieved via ultra-sonification using a Sonoplus HD 2070.2 sonifies with a VS70T probe (Bandelin, Berlin, Germany) at an amplitude of 25% with 1s active and 3 s inactive intervals for 8 min, while the sample was cooled on ice. Cell debris and unbroken cells were collected by centrifugation at 16,000 x *g* for at least 30 min. The supernatant was then filtered through a 0.45 µm pore-size cellulose acetate syringe filter and the cell-free extract was purified via immobilized metal ion chromatography (IMAC) using a 5 mL HisTrap Excel column (Cytiva, Marlborough, MA, United States) attached to a Äkta-Go FPLC device (Cytiva, Marlborough, MA, United States). Elution of column-bound protein was achieved by Imidazole gradient fractionation over 6 column volumes. Collected fractions were analyzed via SDS-PAGE to confirm presence of the target protein. Purified samples were desalted using 15-kDa Amicon centrifugal filters (Merck Millipore, Burlington, MA, United States).

Protein concentration after desalting was determined using the Qubit4 Protein Broad range kit (Thermo Fisher, Waltham, MA, United States).

### Enzyme activity testing

All enzyme assays were carried out at pH 7.4 in 50 mM PBS containing 300 mM NaCl and 0.1% Triton-X. Triton-X, apart from aiding in protein solubility, was used in plastic degradation assays for its surface active properties, ensuring a homogenous dispersion of milled plastic particles in the reaction mixture. To generally assess esterase activity, the purified protein was incubated in cuvettes at a concentration of 1 µM with 0.5 mM of *p*-nitrophenyl butyrate in a total volume of 1 mL; upon ester-bond cleavage, *p*-nitrophenol is released, which was followed at 397 nm.^99^

Polyester depolymerase activity was assessed discontinuously by following the formation of monomers. For determination of PCL-depolymerase activity, 2.5 µM of enzyme was incubated with 5 ± 0.1 mg PCL powder (≤ 100 µm diameter particle size) in 1 mL assay volume in 1.5 mL microcentrifuge tubes. The reactions were incubated at 30 °C with horizontal shaking at 120 rpm for up to 24 h. Sub-samples of 200 μL were taken at multiple timepoints, and the reaction in the sub-samples stopped by addition of 10% (v/v) of 1 M H_2_SO_4_. Acidified samples were centrifuged at maximum speed in a table-top centrifuge and the particle-free supernatant transferred to a HPLC vial.

For determination of LCAP-depolymerase activity, assays were also carried out in 1.5 mL microcentrifuge tubes, in either 0.5 mL or 1 mL total assay volume per reaction. Tubes contained 3 ± 0.1 mg PE-2,18 powder (≤ 100 µm diameter particle size) in 0.5 mL assay volume and purified enzyme was added to a concentration of 2.5 μM. In assays conducted with 1 mL volume, 5 ± 0.1 mg of PE-2,18 powder was used. Samples were incubated at 30°C under horizontal shaking at 120 rpm for up to 72 h. For samples with 1 mL volume, sub-samples of 200 μL were taken at multiple timepoints, and the reactions in the sub-samples were stopped by either addition of an equal volume of isopropanol (200 µl) for samples analyzed by HPLC-MS, or by addition of 1 M H_2_SO_4_ to a concentration of 10% (v/v) for samples analyzed by HPLC-RID. For samples with 0.5 mL volume, the entire sample was used. Assay reactions were conducted in triplicate or in single samples, as specified. Assay buffer with plastic but without enzyme and heat-inactivated enzyme samples supplemented with plastic were included as negative controls. The activity LCPH1 for LCAP degradation was compared directly to that of *Ideonella sakaiensis* PETase (IsPETase),^52^ which was expressed and purified using a pET-21b expression vector kindly provided by the research group of Prof. Uwe Bornscheuer at Universität Greifswald.

Formation of monomer γ-caprolactone was determined with a Shimadzu LC-20 HPLC system (Shimadzu, Kyoto, Japan) equipped with refractive index detector (RID), using 1% (v/v) H_2_SO_4_ in distilled water as mobile phase with a flowrate of 0.6 mL/min and a Rezex™ RHM-Monosaccharide H+ (8%) Ion Exclusion HPLC Column (Phenomenex, Torrance, CA, Unisted States) for separation. LCAP monomer formation was quantified by detection of both PE-2,18-monomers, ethylene glycol and 1,18-octadecanedioate. Ethylene glycol concentrations were determined via HPLC-RID as described for caprolactone.

Octadecanedioate concentrations were determined via HPLC-MS as previously described.^32^ Briefly, an equal volume of MS-grade isopropanol was added to each sample and the samples were mixed and incubated at 37 °C for at least 30 min in order to facilitate dissolution of octadecanedioate and precipitation of protein. Samples were then centrifuged in a table-top centrifuge at maximum speed and the particle-free supernatants were transferred to HPLC vials. Separation and detection of 1,18-octadecanedioate was achieved using MilliQ-water acidified with 0.1% (vol/vol) formic acid and acetonitrile as mobile phases, a Hypersil ODS C18 HPLC column (Thermo Fisher, Waltham, MA, United States), and ESI ionization in the negative mode. All monomers were quantified against authentic standards.

Hydrolysis of β-lactam antibiotics Penicillin G and Ampicillin by GID54916 was determined in 0.5 mL reactions with 500 µM of the respective antibiotic and 2.5 μM purified LCPH1.

The reactions were incubated in glass HPLC vials at room temperature and without shaking in the autosampler of a Shimadzu LC-20 HPLC system (Shimadzu, Kyoto, Japan) equipped with a diode array detector (DAD). Samples were injected at 1 – 2 h intervals for 24 h. Penicillin G and Ampicillin concentrations were determined against authentic standards using a gradient of MilliQ-water with 10 mM ammonium formate (eluent A) and 10% MeOH and (eluent B) 100% MeOH, starting at 45% A at a flow-rate of 0.5 mL/min, and a Nucleosil™ 100-5 C18 column (Machery-Nagel, Düren, Germany); the compounds were detected at 254 nm.

## Data availability

The metagenomic and amplicon sequencing raw data have been deposited at EBI with the identifier PRJEB96049.

## Supporting information

List of candidate plastic depolymerases

## Acknowledgements

The Authors thank Julia Schmidt and Sylke Wiechmann for their excellent technical support, Dr. Nicolai Müller for helpful discussions, and students Diego Casaburi and Asterios Katsioulas for assisting in protein expressions. The authors also thank Prof. Dr. Uwe Bornscheuer for providing the isPETase expression plasmid used in this study.

## Author contributions

**H.L.** and **D.S.** conceived the study. **H.L.** conceived, designed, and performed the experiments, analyzed the data, and interpreted the results. **D.S.** contributed to data interpretation and supervised the project in the capacity of group leader. **N.C.M.** performed SEM analysis and image processing. **M.E.** and **L.B.** synthesized LCAP materials and **M.E.**, **L.B.** and **D.S.** set up the *in-situ* degradation study. **H.L.** and **D.S.** wrote the manuscript. **S.M.** and **D.S.** secured the project funding supporting this research. All authors reviewed and approved the final manuscript.

## References

1. Kibria, Md. G., Masuk, N. I., Safayet, R., Nguyen, H. Q. & Mourshed, M. Plastic waste: challenges and opportunities to mitigate pollution and effective management. Int J Environ Res 17, 20 (2023).

2. da Costa, J. P., Santos, P. S. M., Duarte, A. C. & Rocha-Santos, T. (Nano)plastics in the environment – sources, fates and effects. Science of The Total Environment 566–567, 15–26 (2016).

3. Wright, S. L. & Kelly, F. J. Plastic and human health: a micro issue? Environ Sci Technol 51, 6634–6647 (2017).

4. Marfella, R. et al. Microplastics and nanoplastics in atheromas and cardiovascular events. New England Journal of Medicine 390, 900–910 (2024).

5. Ali, I. et al. Eco-and bio-corona-based microplastics and nanoplastics complexes in the environment: Modulations in the toxicological behavior of plastic particles and factors affecting. Process Safety and Environmental Protection 187, 356–375 (2024).

6. Yang, H. et al. A review of eco-corona formation on micro/nanoplastics and Its effects on stability, bioavailability, and toxicity. Water (Basel) 17, 1124 (2025).

7. Yao, S. et al. Soil metabolome impacts the formation of the eco-corona and adsorption processes on microplastic surfaces. Environ Sci Technol 57, 8139–8148 (2023).

8. Wright, R. J., Langille, M. G. I. & Walker, T. R. Food or just a free ride? A meta-analysis reveals the global diversity of the plastisphere. ISME J 15, 789–806 (2021).

9. Kaur, K. et al. Microplastic-associated pathogens and antimicrobial resistance in environment. Chemosphere 291, 133005 (2022).

10. Joo, S. H., Knauer, K., Su, C. & Toborek, M. Antibiotic resistance in plastisphere. J Environ Chem Eng 13, 115217 (2025).

11. Zhu, D., Ma, J., Li, G., Rillig, M. C. & Zhu, Y.-G. Soil plastispheres as hotspots of antibiotic resistance genes and potential pathogens. ISME J 16, 521–532 (2022).

12. Stevenson, E. M., Buckling, A., Cole, M., Lindeque, P. K. & Murray, A. K. Selection for antimicrobial resistance in the plastisphere. Science of The Total Environment 908, 168234 (2024).

13. Bowley, J., Baker-Austin, C., Porter, A., Hartnell, R. & Lewis, C. Oceanic hitchhikers – assessing pathogen risks from marine microplastic. Trends Microbiol 29, 107–116 (2021).

14. Luo, G., Fan, L., Liang, B., Guo, J. & Gao, S.-H. Determining antimicrobial resistance in the plastisphere: lower risks of nonbiodegradable vs higher risks of biodegradable microplastics. Environ Sci Technol 59, 7722–7735 (2025).

15. Guo, W., Duan, J., Shi, Z., Yu, X. & Shao, Z. Biodegradation of PET by the membrane-anchored PET esterase from the marine bacterium Rhodococcus pyridinivorans P23. Commun Biol 6, 1090 (2023).

16. Puiggené, Ò. et al. Extracellular degradation of a polyurethane oligomer involving outer membrane vesicles and further insights on the degradation of 2,4-diaminotoluene in Pseudomonas capeferrum TDA1. Sci Rep 12, 2666 (2022).

17. Gambarini, V. et al. PlasticDB: a database of microorganisms and proteins linked to plastic biodegradation. Database 2022, baac008 (2022).

18. Buchholz, P. C. F. et al. Plastics degradation by hydrolytic enzymes: the plastics-active enzymes database— PAZy. Proteins: Structure, Function, and Bioinformatics 90, 1443–1456 (2022).

19. Zhang, H. et al. The metagenome-derived esterase PET40 is highly promiscuous and hydrolyses polyethylene terephthalate (PET). FEBS J 291, 70–91 (2024).

20. Wagner, U. G., Petersen, E. I., Schwab, H. & Kratky, C. EstB from Burkholderia gladiolilll: a novel esterase with a β-lactamase fold reveals steric factors to discriminate between esterolytic and β-lactam cleaving activity. Protein Science 11, 467–478 (2002).

21. Müller, C. A. et al. Discovery of polyesterases from moss-associated microorganisms. Appl Environ Microbiol 83, e02641–16 (2017).

22. Tchigvintsev, A. et al. The environment shapes microbial enzymes: five cold-active and salt-resistant carboxylesterases from marine metagenomes. Appl Microbiol Biotechnol 99, 2165– 2178 (2015).

23. Soto-Hernández, A., Muriel-Millán, L. F., Gracia, A., Sánchez-Flores, A. & Pardo-López, L. Enzymatic characterization and polyurethane biodegradation assay of two novel esterases isolated from a polluted river. PLoS One 20, e0327637 (2025).

24. Kwon, S., Yoo, W., Kim, Y.-O., Kim, K. K. & Kim, T. D. Molecular characterization of a novel family VIII esterase with β-lactamase activity (PsEstA) from Paenibacillus sp. Biomolecules 9, 786 (2019).

25. Jeon, J. H. et al. A novel family VIII carboxylesterase hydrolysing third- and fourth-generation cephalosporins. Springerplus 5, 525 (2016).

26. Park, J.-M., Won, S.-M., Kang, C.-H., Park, S. & Yoon, J.-H. Characterization of a novel carboxylesterase belonging to family VIII hydrolyzing β-lactam antibiotics from a compost metagenomic library. Int J Biol Macromol 164, 4650–4661 (2020).

27. Le, L. T. H. L. et al. Dual functional roles of a novel bifunctional β-lactamase/esterase from Lactococcus garvieae. Int J Biol Macromol 206, 203–212 (2022).

28. Jeon, J. H. et al. Novel metagenome-derived carboxylesterase that hydrolyzes β-lactam antibiotics. Appl Environ Microbiol 77, 7830–7836 (2011).

29. Nan, F. et al. A novel family VIII carboxylesterase with high hydrolytic activity against ampicillin from a soil metagenomic library. Mol Biotechnol 61, 892–904 (2019).

30. Mokoena, N., Mathiba, K., Tsekoa, T., Steenkamp, P. & Rashamuse, K. Functional characterisation of a metagenome derived family VIII esterase with a deacetylation activity on β-lactam antibiotics. Biochem Biophys Res Commun 437, 342–348 (2013).

31. Häußler, M., Eck, M., Rothauer, D. & Mecking, S. Closed-loop recycling of polyethylene-like materials. Nature 590, 423–427 (2021).

32. Eck, M., et al. Biodegradable high-density polyethylene-like material. Angewandte Chemie International Edition 62, e202213438 (2023).

33. Nelson, T. F., Rothauer, D., Sander, M. & Mecking, S. Degradable and recyclable polyesters from multiple chain length bio- and waste-sourceable monomers. Angewandte Chemie International Edition 62, e202310729 (2023).

34. Lerner, H., Schleheck, D., Wendlandt, D. & Hupach, S. Unkaputtbar – oder nicht? Nachrichten aus der Chemie 73, 60–62 (2025).

35. Ribitsch, D. et al. Hydrolysis of polyethyleneterephthalate by p -nitrobenzylesterase from Bacillus subtilis. Biotechnol Prog 27, 951–960 (2011).

36. Schneiker, S. et al. Complete genome sequence of the myxobacterium Sorangium cellulosum. Nat Biotechnol 25, 1281–1289 (2007).

37. Costa, O. Y. A. et al. Impact of different trace elements on the growth and proteome of two strains of Granulicella, class Acidobacteriia. Front Microbiol 11, (2020).

38. Teufel, F. et al. SignalP 6.0 predicts all five types of signal peptides using protein language models. Nat Biotechnol 40, 1023–1025 (2022).

39. Zückert, W. R. Secretion of bacterial lipoproteins: through the cytoplasmic membrane, the periplasm and beyond. Biochimica et Biophysica Acta (BBA) - Molecular Cell Research 1843, 1509–1516 (2014).

40. Cea-Rama, I. et al. Crystal structure of a family VIII β-lactamase fold hydrolase reveals the molecular mechanism for its broad substrate scope. FEBS J 289, 6714–6730 (2022).

41. Fernández, I., Bickelhaupt, F. M. & Svatunek, D. Unraveling the Bürgi-Dunitz angle with precision: the power of a two-dimensional energy decomposition analysis. J Chem Theory Comput 19, 7300–7306 (2023).

42. Tanunchai, B. et al. Nitrogen fixing bacteria facilitate microbial biodegradation of a bio-based and biodegradable plastic in soils under ambient and future climatic conditions. Environ Sci Process Impacts 24, 233–241 (2022).

43. Avasthi, I., et al. Biodegradable mineral plastics. Small Methods 8, 2300575 (2024).

44. Meereboer, K. W., Pal, A. K., Cisneros-López, E. O., Misra, M. & Mohanty, A. K. The effect of natural fillers on the marine biodegradation behaviour of poly(3-hydroxybutyrate-co-3-hydroxyvalerate) (PHBV). Sci Rep 11, 911 (2021).

45. Narita, S. & Tokuda, H. An ABC transporter mediating the membrane detachment of bacterial lipoproteins depending on their sorting signals. FEBS Lett 580, 1164–1170 (2006).

46. Wilson, M. M. & Bernstein, H. D. Surface-exposed lipoproteins: an emerging secretion phenomenon in Gram-negative bacteria. Trends Microbiol 24, 198–208 (2016).

47. de Sandozequi, A. & Martínez-Anaya, C. Bacterial surface-exposed lipoproteins and sortase-mediated anchored cell surface proteins in plant infection. Microbiologyopen 12, (2023).

48. El Rayes, J., Rodríguez-Alonso, R. & Collet, J.-F. Lipoproteins in Gram-negative bacteria: new insights into their biogenesis, subcellular targeting and functional roles. Curr Opin Microbiol 61, 25–34 (2021).

49. Huynh, M. S. et al. Reconstitution of surface lipoprotein translocation through the Slam translocon. Elife 11, (2022).

50. Avila-Calderón, E. D. et al. Outer membrane vesicles of Gram-negative bacteria: an outlook on biogenesis. Front Microbiol 12, (2021).

51. Valguarnera, E., Scott, N. E., Azimzadeh, P. & Feldman, M. F. Surface exposure and packing of lipoproteins into outer membrane vesicles are coupled processes in Bacteroides. mSphere 3, (2018).

52. Yoshida, S. et al. A bacterium that degrades and assimilates poly(ethylene terephthalate). Science 351, 1196–9 (2016).

53. Akutsu, Y., Nakajima-Kambe, T., Nomura, N. & Nakahara, T. Purification and properties of a polyester polyurethane-degrading enzyme from Comamonas acidovorans TB-35. Appl Environ Microbiol 64, 62–67 (1998).

54. Doi, R. H., Kosugi, A., Murashima, K., Tamaru, Y. & Han, S. O. Cellulosomes from mesophilic bacteria. J Bacteriol 185, 5907–14 (2003).

55. Salvachúa, D. et al. Outer membrane vesicles catabolize lignin-derived aromatic compounds in Pseudomonas putida KT2440. Proceedings of the National Academy of Sciences 117, 9302– 9310 (2020).

56. Gasser, M. T. et al. Membrane vesicles can contribute to cellulose degradation by Teredinibacter turnerae, a cultivable intracellular endosymbiont of shipworms. Microb Biotechnol 17, e70064 (2024).

57. Rudnicka, M. et al. Outer membrane vesicles as mediators of plant–bacterial interactions. Front Microbiol 13, (2022).

58. Chen, Z. et al. Biodegradation of highly crystallized poly(ethylene terephthalate) through cell surface codisplay of bacterial PETase and hydrophobin. Nat Commun 13, 7138 (2022).

59. Ercoli, G. et al. LytM proteins play a crucial role in cell Separation, outer membrane composition, and pathogenesis in nontypeable Haemophilus influenzae. mBio 6, (2015).

60. Park, S. & Sauer, K. Controlling biofilm development through cyclic di-GMP signaling. Adv Exp Med Biol 1386, 69–94 (2022).

61. Hu, X.-M. et al. Bacterial c-di-GMP signaling gene affects mussel larval metamorphosis through outer membrane vesicles and lipopolysaccharides. NPJ Biofilms Microbiomes 10, 38 (2024).

62. Wilson, M. M., Anderson, D. E. & Bernstein, H. D. Analysis of the outer Membrane proteome and secretome of bacteroides fragilis reveals a multiplicity of secretion mechanisms. PLoS One 10, e0117732 (2015).

63. Dhurve, G., Madikonda, A. K., Jagannadham, M. V. & Siddavattam, D. Outer membrane vesicles of Acinetobacter baumannii DS002 are selectively enriched with TonB-dependent transporters and play a key role in iron acquisition. Microbiol Spectr 10, (2022).

64. Eck, M. Degradable polyethylene-like materials. (Universität Konstanz, Konstanz, 2023).

65. Lott, C. et al. Field and mesocosm methods to test biodegradable plastic film under marine conditions. PLoS One 15, e0236579 (2020).

66. Larsen, M. H., Biermann, K., Tandberg, S., Hsu, T. & Jacobs William R., Jr. Genetic manipulation of Mycobacterium tuberculosis. Curr Protoc Microbiol 6, 10A.2.1–10A.2.21 (2007).

67. Berthelet, M., Whyte, L. G. & Greer, C. W. Rapid, direct extraction of DNA from soils for PCR analysis using polyvinylpolypyrrolidone spin columns. FEMS Microbiol Lett 138, 17–22 (1996).

68. Klindworth, A. et al. Evaluation of general 16S ribosomal RNA gene PCR primers for classical and next-generation sequencing-based diversity studies. Nucleic Acids Res 41, 1–11 (2013).

69. Purahong, W. et al. Back to the future: decomposability of a biobased and biodegradable plastic in field soil environments and Its microbiome under ambient and future climates. Environ Sci Technol 55, 12337–12351 (2021).

70. Lerner, H. et al. Culture-independent analysis of linuron-mineralizing microbiota and functions in on-farm biopurification systems via DNA-Stable Isotope Probing: comparison with enrichment culture. Environ Sci Technol 54, 9387–9397 (2020).

71. Quast, C. et al. The SILVA ribosomal RNA gene database project: Improved data processing and web-based tools. Nucleic Acids Res 41, 590–596 (2013).

72. Abarenkov, K. et al. The UNITE database for molecular identification and taxonomic communication of fungi and other eukaryotes: sequences, taxa and classifications reconsidered. Nucleic Acids Res 52, D791–D797 (2024).

73. McMurdie, P. J. & Holmes, S. phyloseq: An R Package for Reproducible Interactive Analysis and Graphics of Microbiome Census Data. PLoS One 8, e61217 (2013).

74. Pagès, H., Aboyoun, P., Gentleman, R. & DebRoy, S. Biostrings: Efficient manipulation of biological strings. 10.18129/B9.bioc.Biostrings (2019).

75. Paradis, E. & Schliep, K. ape 5.0: an environment for modern phylogenetics and evolutionary analyses in R. Bioinformatics 35, 526–528 (2019).

76. Oksanen J, Simpson G, Blanchet F, Kindt R, L. P., et al. vegan: Community Ecology Package. https://cran.r-project.org/package=vegan (2024).

77. Lin, H. & Peddada, S. Das. Analysis of compositions of microbiomes with bias correction. Nat Commun 11, 3514 (2020).

78. Wickham, H. Ggplot2: Elegant Graphics for Data Analysis. (Springer-verlag, New York, 2016).

79. Eren, A. M., Vineis, J. H., Morrison, H. G. & Sogin, M. L. A filtering method to generate high quality short reads using Illumina paired-end technology. PLoS One 8, e66643 (2013).

80. Eren, A. M. et al. Community-led, integrated, reproducible multi-omics with anvi’o. Nat Microbiol 6, 3–6 (2020).

81. Langmead, B. & Salzberg, S. L. Fast gapped-read alignment with Bowtie 2. Nat Methods 9, 357–359 (2012).

82. Hyatt, D. et al. Prodigal: prokaryotic gene recognition and translation initiation site identification. BMC Bioinformatics 11, 119 (2010).

83. Parks, D. H. et al. GTDB: an ongoing census of bacterial and archaeal diversity through a phylogenetically consistent, rank normalized and complete genome-based taxonomy. Nucleic Acids Res 50, D785–D794 (2022).

84. Danso, D. et al. New Insights into the Function and Global Distribution of Polyethylene Terephthalate (PET)-Degrading Bacteria and Enzymes in Marine and Terrestrial Metagenomes. Appl Environ Microbiol 84, (2018).

85. Katoh, K. & Standley, D. M. MAFFT multiple sequence alignment software version 7: improvements in performance and usability. Mol Biol Evol 30, 772–780 (2013).

86. Eddy, S. R. Accelerated Profile HMM Searches. PLoS Comput Biol 7, e1002195 (2011).

87. Camacho, C., et al. BLAST+: architecture and applications. BMC Bioinformatics 10, 421 (2009).

88. Lê, S., Josse, J. & Husson, F. FactoMineRlll: an R package for multivariate analysis. J Stat Softw 25, 1–18 (2008).

89. Abramson, J. et al. Accurate structure prediction of biomolecular interactions with AlphaFold 3. Nature 630, 493–500 (2024).

90. Laskowski, R. A., Watson, J. D. & Thornton, J. M. ProFunc: a server for predicting protein function from 3D structure. Nucleic Acids Res 33, W89–W93 (2005).

91. Weininger, D. SMILES, a chemical language and information system. 1. Introduction to methodology and encoding rules. J Chem Inf Comput Sci 28, 31–36 (1988).

92. O’Boyle, N. M. et al. Open Babel: an open chemical toolbox. J Cheminform 3, 33 (2011).

93. Morris, G. M. et al. AutoDock4 and AutoDockTools4: automated docking with selective receptor flexibility. J Comput Chem 30, 2785–2791 (2009).

94. Goddard, T. D. et al. UCSF ChimeraX: meeting modern challenges in visualization and analysis. Protein Science 27, 14–25 (2018).

95. Katoh, K., Rozewicki, J. & Yamada, K. D. MAFFT online service: multiple sequence alignment, interactive sequence choice and visualization. Brief Bioinform 20, 1160–1166 (2019).

96. Hunter, J. D. Matplotlib: a 2D graphics environment. Comput Sci Eng 9, 90–95 (2007).

97. Cock, P. J. A. et al. Biopython: freely available Python tools for computational molecular biology and bioinformatics. Bioinformatics 25, 1422–1423 (2009).

98. Novy, R. & Morris, B. Use of glucose to control basal expression in the pET system. Innovations 13, 13–15 (2001).

99. Peng, Y., Fu, S., Liu, H. & Lucia, L. A. Accurately determining esterase activity via the isosbestic point of p-nitrophenol. Bioresources 11, (2016).

