## Supporting Information for "A Family VIII Esterase with Dual Activities: Bioplastic Depolymerization and Beta-Lactam Antibiotics Hydrolysis"

Supplementary information, includes: Supplementary tables 1-2, Supplementary figures 1-11

**Table S1 Analysis of utilized Forest soil according to Albrecht Standard** conducted by Geobüro Christophel (Lauterhofen, Germany).

| Parameter | Value | Units |
|---|---|---|
| **Cations** | | |
| Potassium | 179 | kg/ha |
| Sodium | 28 | kg/ha |
| Calcium | 1193 | kg/ha |
| Magnesium | 374 | kg/ha |
| **Anions** | | |
| Sulfur | 3 | ppm |
| Phosphorus (available) | 3.1 | kg/ha |
| Phosphorus (stored) | 151 | kg/ha |
| **Trace Elements** | | |
| Boron | 0.6 | ppm |
| Iron | 794.4 | ppm |
| Manganese | 103.3 | ppm |
| Copper | 1.1 | ppm |
| Zinc | 16.8 | ppm |
| **General Soil Data** | | |
| pH ($H_2O$) | 4.3 | |
| Total CEC* | 13.3 | mmol/100g |
| Humus Content | 15.4 | % |
| Total Nitrogen | 0.52 | % |
| C/N Ratio | 17.2 | |

*Cation exchange capacity

**Table S2 concentration of trace minerals and salts added to soil microcosms after 66 days.** The listed concentrations refer to the total water content of the soil based on determined water content and additionally supplied water including the nutrient solution.

| ***Trace Minerals and Salts*** | ***Concentration [µM]*** |
|---|---|
| $Na_2EDTA \cdot 2H_2O$ | 13.40 |
| $FeCl_2 \cdot 4H_2O$ | 7.20 |
| $ZnCl_2$ | 0.35 |
| $MnCl_2 \cdot 4H_2O$ | 0.15 |
| $H_3BO_3$ | 4.85 |
| $CoCl_2 \cdot 6H_2O$ | 0.84 |
| $CuCl_2 \cdot 2H_2O$ | 0.06 |
| $NiCl_2 \cdot 6H_2O$ | 0.08 |
| $Na_2MoO_4 \cdot 2H_2O$ | 0.12 |
| $CaCl_2 \cdot 2H_2O$ | 6.80 |

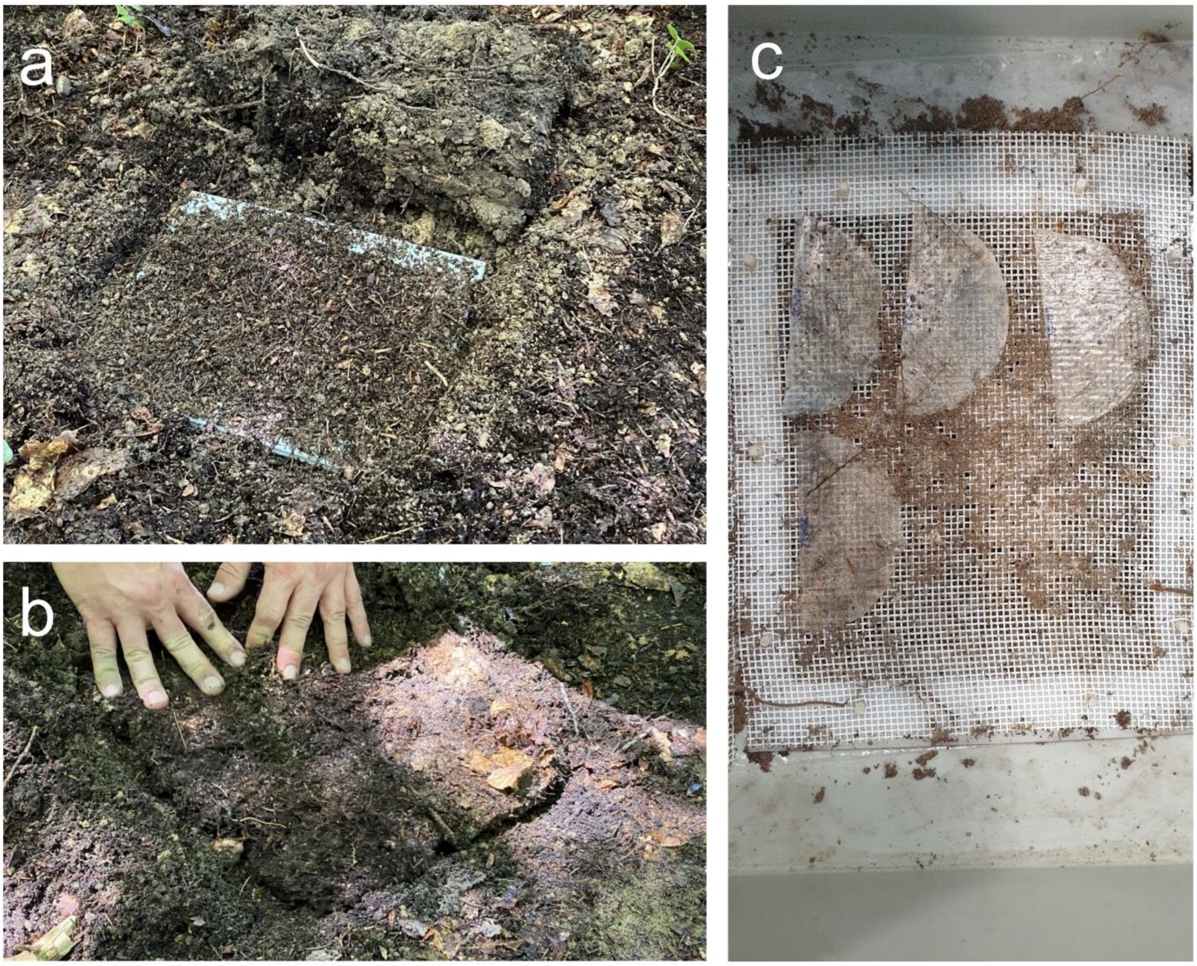


**Figure S1.** Photographs illustrating the placement of the LCAP films mounted into 'HYDRA test frames' (see Lott et al.[1]) under a layer of sieved humus (a) and forest top-soil (b), and of the macroscopical appearance of the LCAP films after recovery and opening of the test frames (c). For scanning electron microscopy (SEM) images of the surfaces of the plastic films before and after incubation, see Fig. S2 and in the main text Fig. 1.

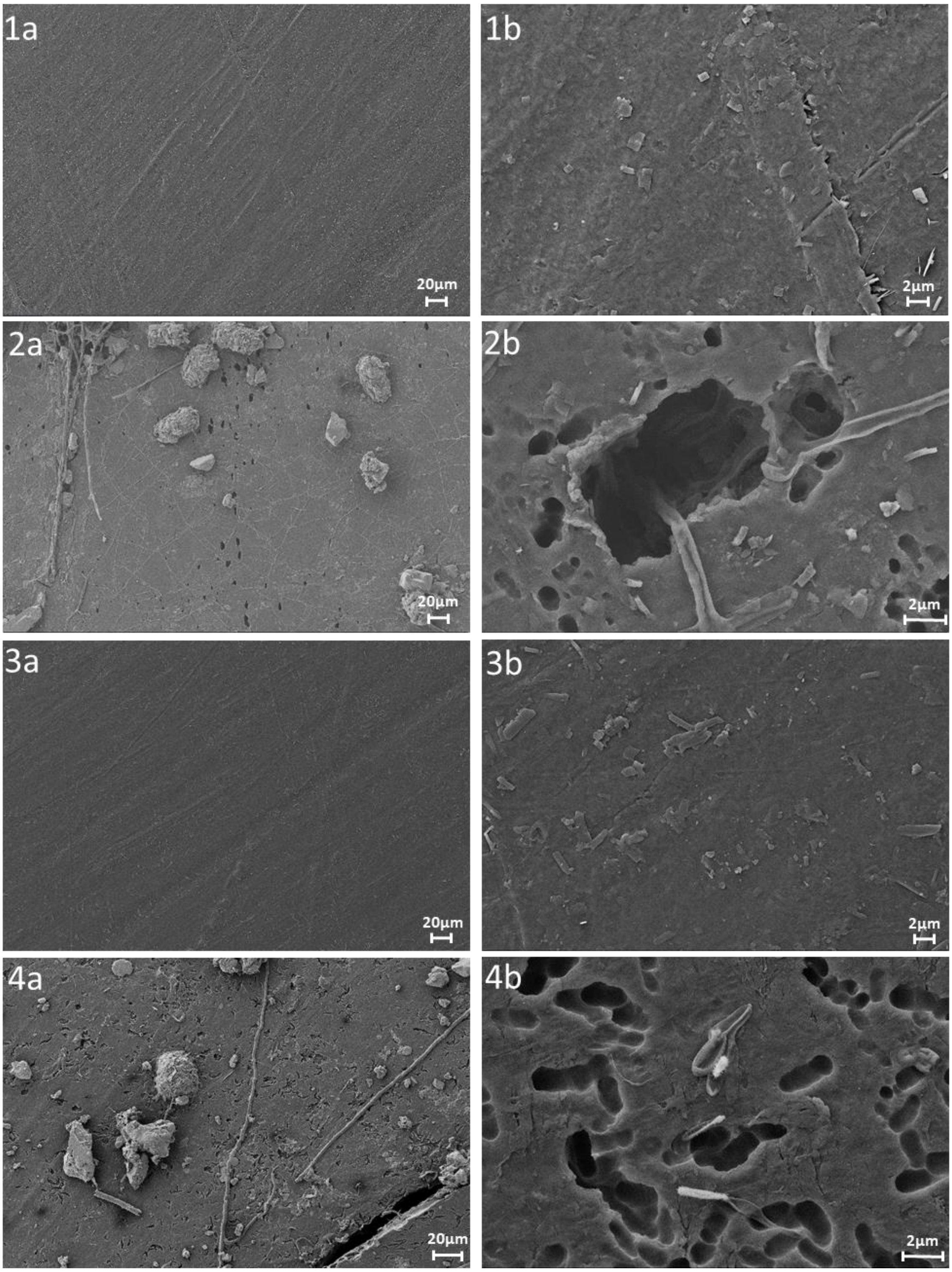


**Figure S2. Scanning electron microscopy (SEM) images of PE-12,12 and PE-18,18 films incubated in pristine forest soil for about one year.** The images were taken at the same position with two different magnifications, as indicated by 20-µm and 2-µm scale bars. The structural damage on the PE-12,12 and PE-18,18 films after incubation (images 2a and 2b, and 4a and 4c, respectively), which was absent for non-incubated foils (images 1a and 1b, and 3a and 3b, respectively), most likely resulted from microbial degradation of the plastic material. See also the additional images in the main text (Fig. 1).

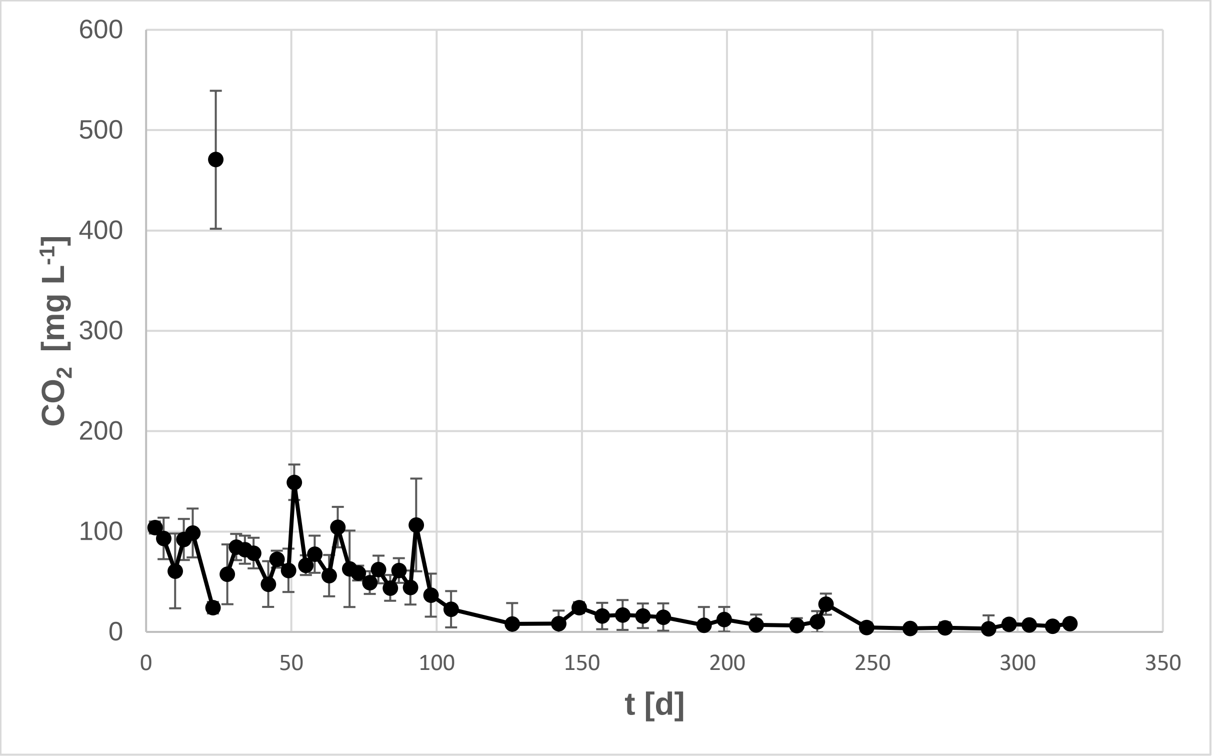


**Figure S3 Average daily $CO_2$ rates in soil blank microcosms.** Rates were determined by division of the measured $CO_2$ concentration at a given time point by the amount of days since the previous measurement. Error bars show the standard deviation of the triplicate microcosms. The outlier observed at 24 d relates to the observed saturation of the NaOH solution at the previous time point, likely resulting in accumulation of $CO_2$ in the microcosm headspaces.

a

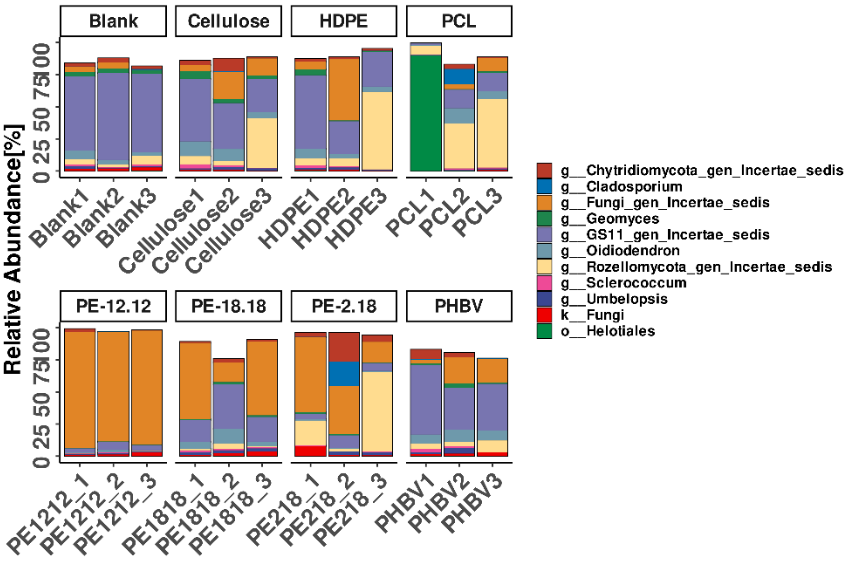


b

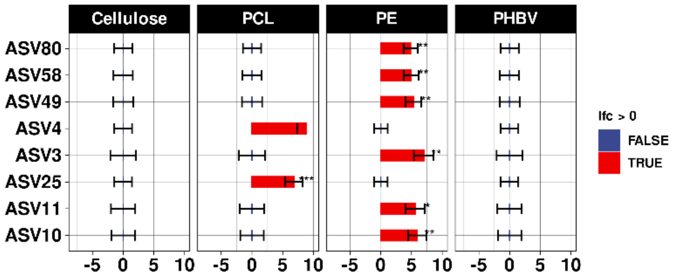


**Figure S4. Bar plots illustrating the ITS gene diversity (genus level) of the fungal communities in soil samples incubated with the different polymers and Log-fold change (LFC) analysis for differentially abundant taxa. (a)** The community composition is shown for all replicate treatments. ASVs assigned to the same genus were combined. When a genus could not be determined with sufficient confidence, the last taxonomic level assigned is indicated. Only ASV groups with a mean relative abundance of >1% are depicted. **(b)** Non-spiked microcosms (blanks) and the HDPE-spiked microcosms were grouped together as reference. Likewise, all microcosms treated with the three types of long-chain PEs were grouped together, while replicates of the cellulose-, PCL- and PHBV-treatments formed separate groups. To reduce the impact of sequencing artifacts and extremely rare ASVs, only ASVs with a count of ≥100 across 10% of samples were included in the analysis. Red bars indicate a significant increase in a taxon’s abundance compared to the control condition. Blue bars indicate a reduction. Statistical significance of the LFC is indicated next to the bars (*:  P ≤ 0.05, **: P ≤ 0.01, ***: P ≤ 0.001, ****: P ≤ 0.0001).

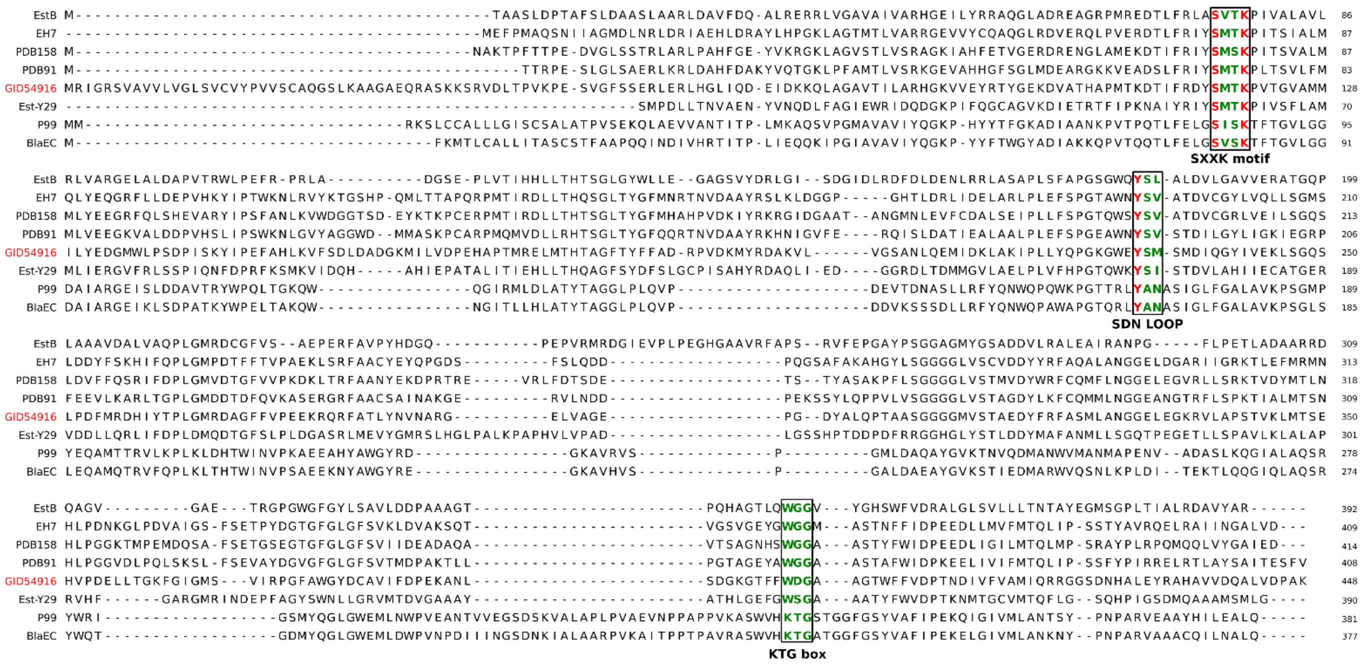


**Fig S5. Multiple sequence alignment of family VIII esterases** EstB from *Burkholderia gladioli*, Est-14 (GeneBank KY483644), the PlasticDB (PDB) entries 91 and 158 and class C β-lactamases p99 from *Enterobacter cloacae*, BlaEC from *Escherichia coli* and the metagenomic penicillin-binding beta-lactamase homolog Est-Y29, in comparison to GID54916 (red). The key conserved regions are highlighted in green, while residues making up the catalytic triad are highlighted in red.

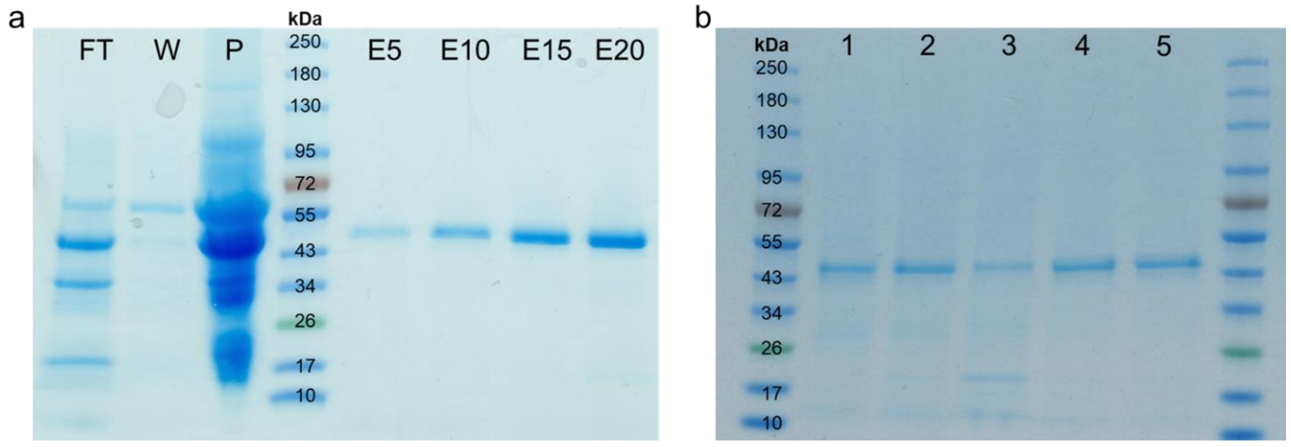


**Fig S6 SDS-PAGE of purified GID54916 expressions used in this study**. (a) Exemplary expression showing the different fractions collected during the purification process. His-Trap column flow-through (FT), the collected wash fraction (W), the proteins present in the cell pellet (P) and the eluted (E) protein applied in different volumes (5, 10, 15 and 20 µL). (b) SDS-PAGE of the different utilized purified GID54916 expressions at working concentration (2.5 µM). The estimated size of GID54916, including His-Tag, is 46.3 kDa. No, or only minor, contaminated binds are present in the purified preparations.

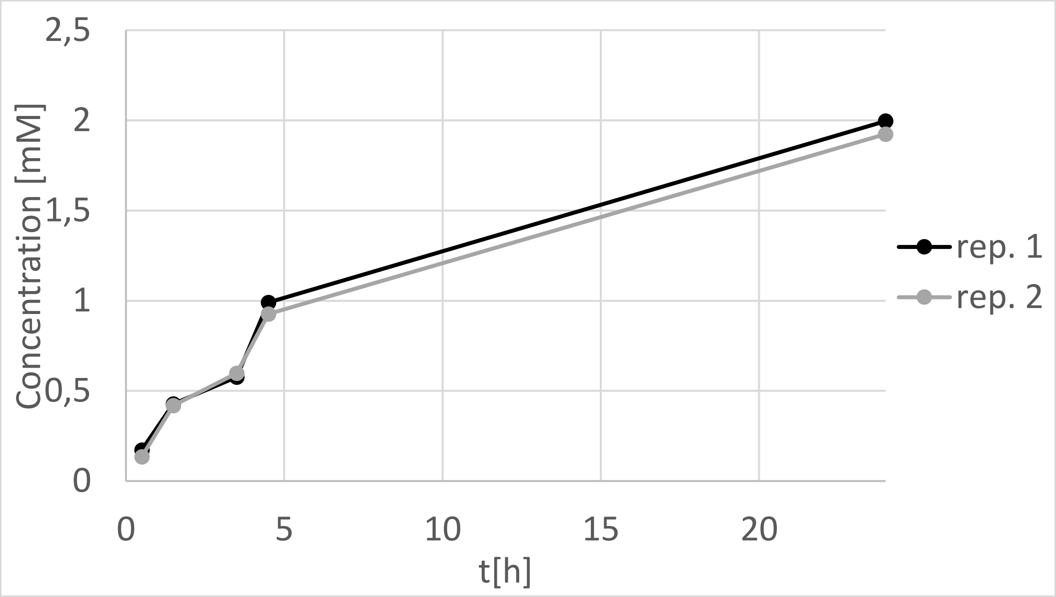


**Fig S8. Time course of ε-caprolactone production over 24 hours by GID54916 from PCL powder.** Reactions contained 5 mg of PCL and 2.5 µM enzyme in 1mL PBS Buffer with 0.1% Triton-X. Reactions were set up in duplicate and samples of 200 µL taken at 5 time points over 24h. Reactions were stopped with 10% (v/v) of 1 M $H_2SO_4$ and ε-caprolactone detected via HPLC-RID.

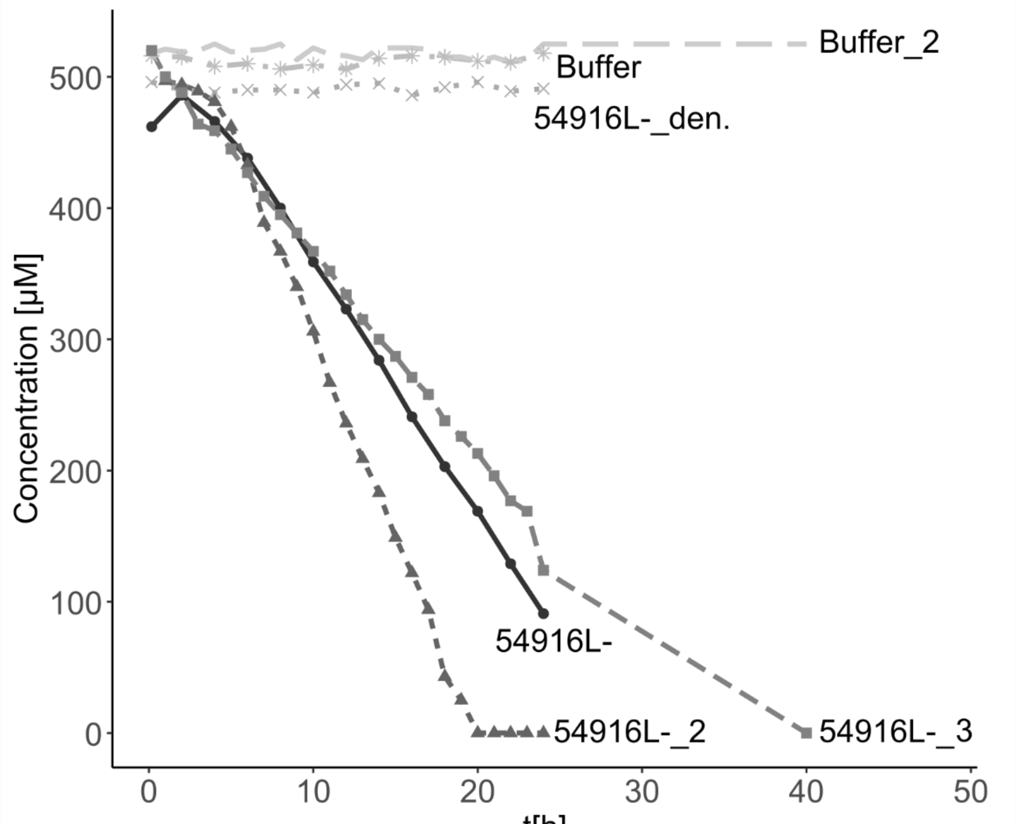


**Fig S9. Time course of Penicillin G hydrolysis by GID54916 over 24-40 hours, measured as declining Penicillin G concentration by HPLC-DAD.** Reactions contained 500 µM Penicillin G and 2.5 µM GID54916 from three separate expressions (numbered), heat-denatured GID54916, or a PBS buffer control (0.05% Triton-X) in 0.5 mL total volume. Samples were taken every 1-2 hours for 24 hours. In case of expression 3 and Buffer 2, an additional final measurement was conducted after 40h to assure complete substrate depletion.

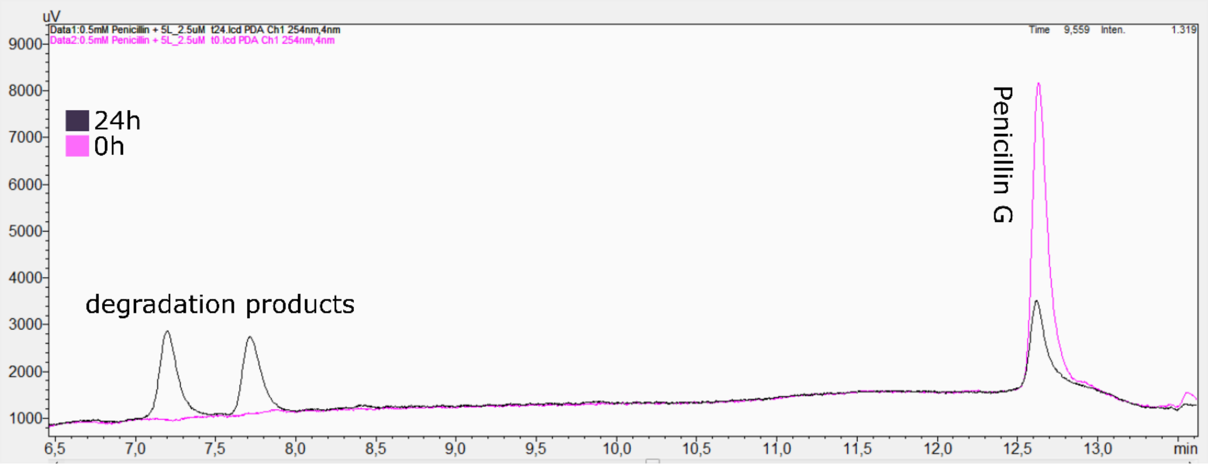


**Fig S10. Exemplary HPLC chromatogram of Penicillin G diminishment and formation of degradation product in response to 24h of GID54916 exposure.** Penicillin G peak was observed at ~12.7 min at initiation of the reaction (pink) and a residual peak is visible after 24 h (black). After 24 h of incubation two additional peaks were observed at 7.2 and 7.7 min, likely representing penicilloic acid and a second intermediary degradation product, respectively.

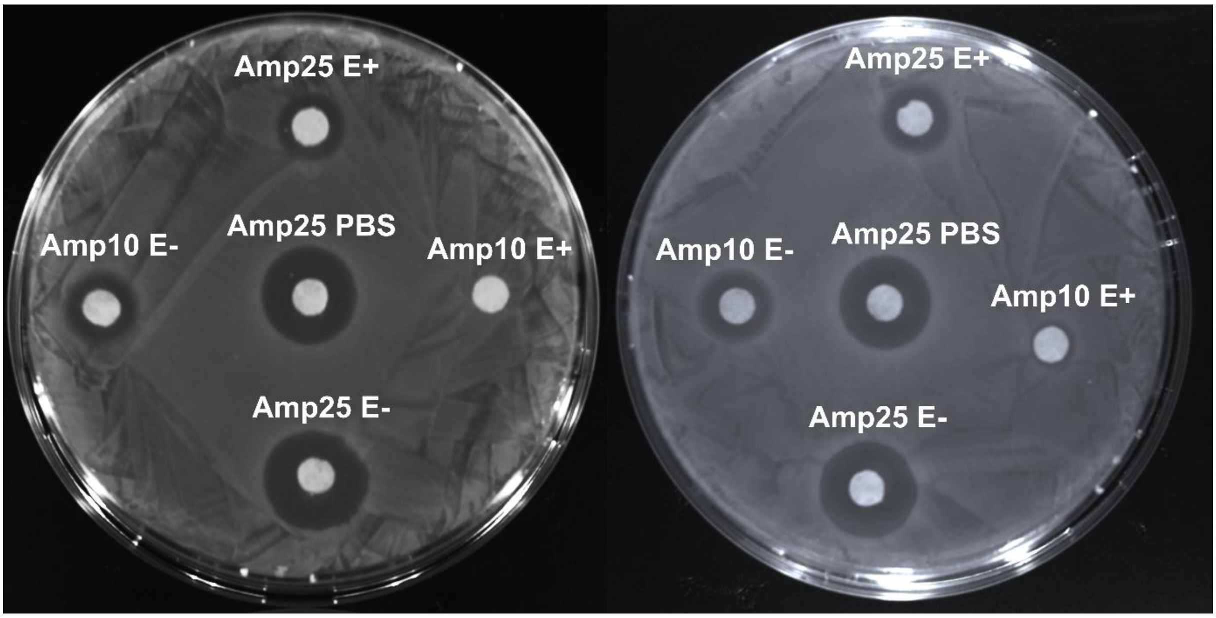


**Fig S11. Ampicillin disc diffusion assay with and without GID54916 exposure in duplicate.** Filter paper discs supplemented with 10 or 25 µg of Ampicillin were incubated over night at 30°C with 20 µL of 5 µM GID54916 in PBS buffer (E+), 5 µM GID54916 after heat-denaturation (E-) or PBS buffer without enzyme (PBS). The discs were placed on Mueller–Hinton agar plates previously streaked with Ampicillin susceptible *E. coli* K12 cells. Plates were incubated over night at 37°C and plates photographed. Inhibition zones are clearly diminished when discs were exposed to active GID54916 for both Ampicillin concentrations.

### References

1. Lott, C. *et al.* Field and mesocosm methods to test biodegradable plastic film under marine conditions. *PLoS One* **15**, e0236579 (2020).
